# An efficient analytical reduction of detailed nonlinear neuron models

**DOI:** 10.1101/506485

**Authors:** Oren Amsalem, Guy Eyal, Noa Rogozinski, Felix Schürmann, Michael Gevaert, Pramod Kumbhar, Idan Segev

**Author notes:** Correspondence: Oren Amsalem.

## Abstract

Detailed conductance-based nonlinear neuron models consisting of thousands of synapses are key for understanding of the computational properties of single neurons and large neuronal networks, and for interpreting experimental results. Simulations of these models are computationally expensive, considerably curtailing their utility. *Neuron_Reduce* is a new analytical approach to reduce the morphological complexity and computational time of nonlinear neuron models. Synapses and active membrane channels are mapped to the reduced model preserving their transfer impedance to the soma; synapses with identical transfer impedance are merged into one NEURON process still retaining their individual activation times. *Neuron_Reduce* accelerates the simulations by 40-250 folds for a variety of cell types and realistic number (10,000-100,000) of synapses while closely replicating voltage dynamics and specific dendritic computations. The reduced neuron-models will enable realistic simulations of neural networks at unprecedented scale, including networks emerging from micro-connectomics efforts and biologically-inspired “deep networks”. *Neuron_Reduce* is publicly available and is straightforward to implement.

## Introduction

Compartmental models (CMs) were first employed by Wilfrid Rall to study the integrative properties of neurons^1^. They enabled him to explore the impact of spatio-temporal activation of conductance-based dendritic synapses on the neuron’s output and the effect of the dendritic location of a synapse on the time course of the somatic excitatory post-synaptic potential^2^. By simulating electrically-distributed neuron models, Rall demonstrated how the cable properties of dendrites explain the variety of somatic EPSP shapes that were recorded at the soma of α-motoneurons, thus negating the dominant explanation at that time that the differences in shapes of the somatic EPSPs in these cells result from differences in the kinetics of the respective synapses. This was an impressive example that faithful models of the neuron (as a distributed rather than a “point” electrical unit) are essential for the correct interpretation of experimental results. Since Rall’s 1964 and 1967 studies using CMs, EPSP “shape indices” measured at the soma are routinely used for estimating the electrotonic distance of dendritic synapses from the soma.

Over the years, detailed CMs of neurons have provided key insights into hundreds of experimental findings, both at the single cell and the network levels. A notable example at the single cell level is the explanation as to why the somatic *Na^+^* action potential propagates backward in the soma-to-dendrites direction and (typically) not *vice versa*^3^. CMs have also pinpointed the conditions for the generation of local dendritic *Ca^2+^* spikes^4–6^ and provided an explanation for the spatial restriction of the active spread of dendritic spikes from distal dendrites to the soma^7^ and see also^8–14^. Today, detailed CMs models are even been used for simulating signal processing in human pyramidal neurons, including their large numbers of dendritic spines/synapses^15^.

At the network level, detailed CMs are utilized for such noteworthy projects as large-scale simulations of densely *in silico* reconstructed cortical circuits^16, 17^ and the overarching goal of the Allen Institute to simulate large parts of the visual system of the mouse^18, 19^. Because detailed compartmental modeling is increasingly becoming an essential tool for understanding diverse neuronal phenomena, major efforts have been invested in developing user-friendly computer software that implements detailed CMs, the best known of which are NEURON^20^, GENESIS^21^, NeuroConstruct^22^, PyNN^23^ and recently BioNet^24^, NTS^25^, NetPyNE^26^ and Geppetto^27^.

Modern personal computers can simulate tens of seconds of electrical activity of single neurons comprising thousands of nonlinear compartments and synapses. However, they handle poorly cases where many model configurations need to be evaluated such as in large-scale parameter fitting for single-neuron models^5, 28^, or when the dendritic tree is morphologically and electrically highly intricate and consists of tens of thousands of dendritic synapses, as with cortical pyramidal neurons^15^. When the aim is to simulate a neuronal network consisting of hundreds of thousands of such neurons, only very powerful computers can cope. For example, the simulation of a cortical network consisting of 200,000 detailed neuron models on the BlueGene/Q supercomputer takes several hours to simulate 30 seconds of biological time^17^.

To overcome this obstacle, two approaches have been pursued. The first involves developing alternative, cheaper and more efficient computing architectures (e.g., neuromorphic-based computers^29, 30^). These have not yet reached the stage where they can simulate large-scale network models with neurons consisting of branched nonlinear dendrites having a realistic number of synapses. The other approach is to simplify neuron models while preserving their input/output relationship as faithfully as possible. Rall^31^ was the first to suggest a reduction scheme in his “equivalent cylinder” model, which showed that, for certain idealized passive dendritic trees, the whole tree could be collapsed into a single cylinder that was analytically identical to the detailed tree. The “equivalent cylinder” preserves the total dendritic membrane area, the electrotonic length of the dendrites, and most importantly, the post-synaptic potential (amplitude and time-course) at the soma for a dendritic synapse when mapped to its respective electrotonic location on the “equivalent cylinder”^32, 33^. However, this method is not applicable for dendritic trees with large variability in their cable lengths (e.g., pyramidal neurons with a long apical tree and short basal trees), conductance-based synapses, or for dendrites with nonlinear membrane properties.

Over the years, several different reduction schemes have been proposed; e.g., a recent work mapped all the synapses to a single compartment, taking the filtering effect of the dendrites into account^34^. Other methods reduce the detailed morphology to a simplified geometric model while preserving the total membrane area^35–37^ or the axial resistivity^38^; see also^12, 39, 40^. However, these methods have a variety of drawbacks; in particular, they are either “hand fitted” and thus lack a clear analytical underpinning or are complicated to implement, and in some cases, their computational advantage for realistic numbers (thousands) of synapses is not quantified. Most of these methods do not support dendrites with active conductances^35, 38, 39, 41, 42^ and they have not been tested on a broad range of neuron types. Importantly, none of the previous methods provided an easy-to-use open access implementation. Thus, today there is no simple, publicly available reduction method for neuron models that can be used by the extensive neuroscience and machine-learning communities.

To respond to this need, the present study provides an analytic method for reducing the complexity of detailed neuron models while faithfully preserving the essential input/output properties of these models. *Neuron_Reduce* is based on key theoretical insights from Rall’s cable theory, and its implementation for any neuron type is straightforward without requiring hand tuning. Depending on the neuron modeled and the number of synapses, *Neuron_Reduce* accelerates the simulation run-time by a factor of up to 250 while preserving the identity of individual synapses and their respective dendrites. It also preserves specific membrane properties and dendritic nonlinearities, hence maintaining specific dendritic computations. *Neuron_Reduce* is easy to use, fully documented, and publicly available on GitHub (https://github.com/orena1/neuron_reduce).

## Results

### Analytic mapping of a detailed dendritic tree to a reduced multi-cylinder tree

The thrust of our analytical reduction method (*Neuron_Reduce*) is described in **Fig. 1a-c**. This method is based on representing each of the original stem dendrites by a single cylindrical cable which has the same specific membrane resistivity (*Rm,* in Ωcm^2^), capacitance (*Cm*, in F/cm^2^) and axial resistivity (*Ra*, in Ωcm) as in the detailed tree (**Fig. 1a**). Also, each cylindrical cable satisfies two constraints: (i) the magnitude of the transfer impedance, |*Z*_0,*L*_(*ω*)| = |*V*_0_(*ω*)/*I_L_*(*ω*)|, from its distal sealed end (*X = L*) to its origin at the soma-end (*X = 0*) is identical to the magnitude of the transfer impedance from the electrotonically most distal dendritic tip to the soma in the respective original dendrite; (ii) at its proximal end (X = 0), the magnitude of the input impedance, |*Z*_0,0_(*ω*)| = |*V*_0_(*ω*)/*I_0_*(*ω*)|, is identical to that of the respective stem dendrite (when decoupled from the soma). As shown in Eqs. (1) – (11) (**Methods**), these two constraints, while preserving the specific membrane and axial properties, guarantee a unique cylindrical cable (with a specific diameter and length) for each of the original dendrites.

**Figure 1.**
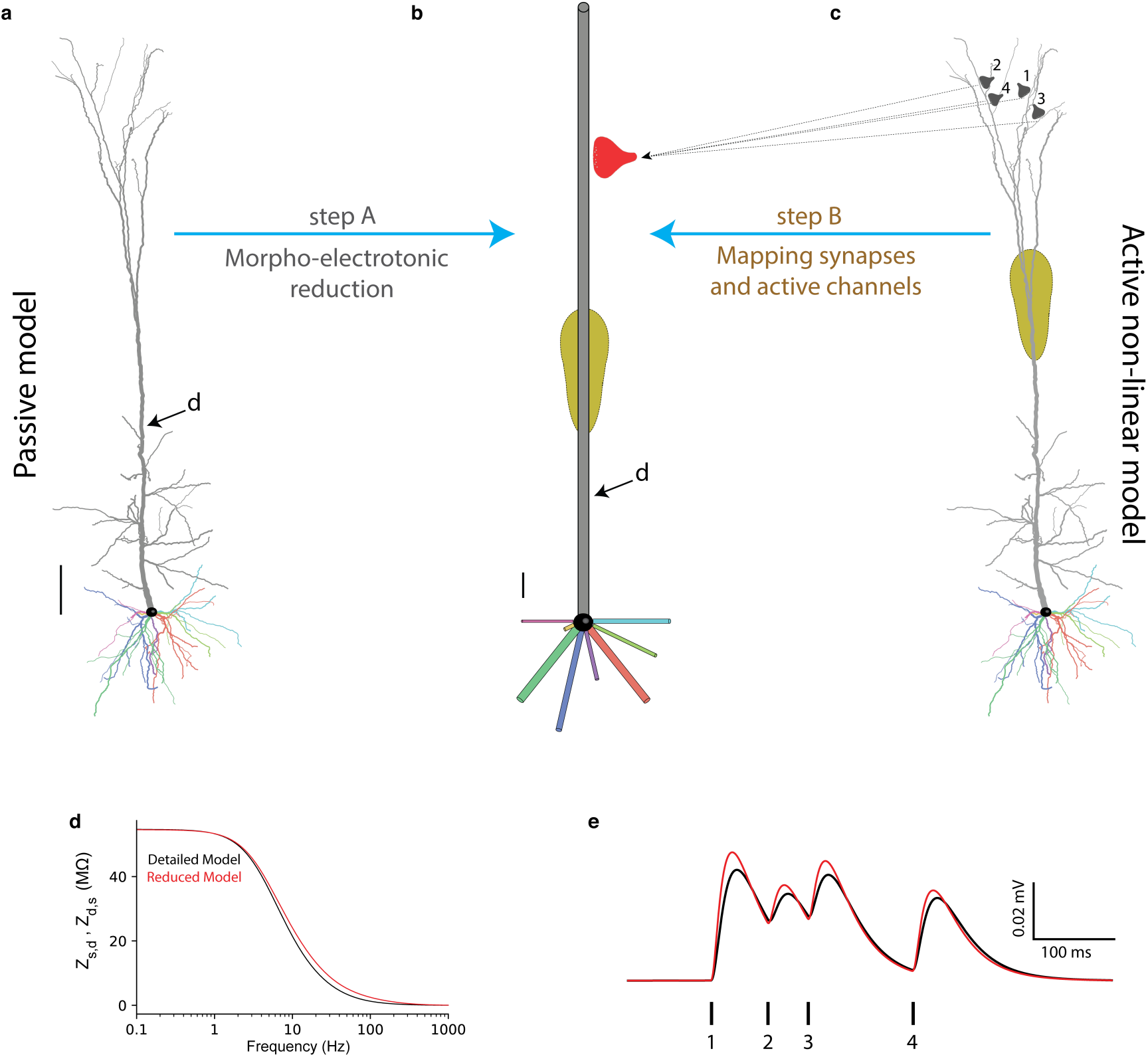
An analytic method for reducing neuron model complexity (*Neuron_Reduce*). **a** Detailed passive model of 3D reconstructed L5 thick tufted pyramidal cell from rat neocortex. Its nine stem dendrites (one apical and 8 basal) are depicted in different colors. **b** Each original stem dendrite is reduced to a single cylinder that retains the specific passive cable properties (*Rm*, *Cm*, and *Ra*) of the original tree. The diameter and length of the respective cylinders are computed analytically using Eqs. (1) – (11), such that each cylinder preserves both the transfer resistance from the most electrotonically distal dendritic tip to the soma as well as the input resistance at the soma end of the corresponding stem dendrite. This generates a unique cylindrical cable for each of the original stem dendrites. Scale bars in a and b are 100 µm. **c** Synapses with similar transfer resistance to the soma (exemplar synapses are marked as 1-4 at top right) are all mapped to the respective locus in the reduced cylinder so that their transfer resistance is similar in the two models. In the reduced model, these synapses are merged into one “NEURON” process (red synapse in b), but they retain their individual activation time (see **Methods** and Supplementary Fig. 1). The same mapping also holds for active membrane conductances (schematic yellow region, denoting the *Ca^2+^* “hot spot” in the apical tree). **d** Transfer impedance (*Z_d_*_,0_ = *Z*_0,*d*_) between point d on the apical tree (shown in a and b) and the soma (X = 0) as a function of the input frequency in both the detailed (black trace) and the reduced (red trace) models. **e** Composite somatic EPSPs resulting from sequential activation of the four distal apical synapses shown in c in the detailed model (black trace) and in the reduced model (red trace). In this simulation the dendritic tree was passive. The synapses were activated in temporal order 1, 2, 3, 4 as shown by the vertical lines below the composite EPSP. The respective peak conductances of these AMPA-based synapses were 0.6, 0.3, 0.4 and 0.4 nS (see details in Supplementary Table 2 and see Supplementary Fig. 1 for the active case).

Because the transfer impedance in both the original dendrite and in the respective cylindrical cable spans from |*Z*_0,*L*_(*ω*)| to |*Z*_0,0_(*ω*)|, all dendritic loci having intermediate transfer impedance values can be mapped to a specific locus in the respective cylinder that preserves this intermediate transfer impedance. This mapping guarantees (for the passive case) that the magnitude of the somatic voltage response, *V*_0_(*ω*), to an input current, *I_x_*(*ω*), injected at a dendritic location, *x*, will be identical in both the detailed and the reduced cylinder models (see **Methods**). Consequently, synapses and nonlinear ion channels are mapped to their respective loci in the reduced cylinder while preserving the respective transfer resistance to the soma (see **Fig. 1** Step B and **Methods**). Based on Eqs. (1) – (11), *Neuron_Reduce* generates a reduced multi-cylindrical tree for any ω value (different reduced models for different ω values). Conveniently, we found a close match between the detailed and the reduced models for ω = 0 (the steady-state case); Therefore, all figures in this work are based on reduced models with ω = 0 (see **Discussion**)

### *Neuron_Reduce* implemented on layer 5 pyramidal cell model with synapses

In **Figure 1**, *Neuron_Reduce* is implemented on a detailed compartmental model of a 3D reconstructed layer 5 pyramidal neuron from the rat somatosensory cortex (same model as in ^5^). This neuron consists of eight basal dendrites and one apical dendrite (shown in different colors) stemming from the soma. This neuron model has active membrane ion channels at both the soma and dendrites (see below). However, *Neuron_Reduce* first treats the modeled tree as passive by abolishing all voltage-dependent membrane conductances, and only retaining the leak conductance. Implementing Eqs. (1) - (11) for this cell produced a reduced, multi-cylindrical, passive model (**Fig. 1b**, Step A) consisting of only 50 compartments rather than the 642 compartments in the detailed model.

**Figure 1c** shows an example of four synapses located at different apical branches. These synapses all have the same transfer resistance to the soma in the detailed tree. Therefore, *Neuron_Reduce* maps these synapses to a single respective locus in the respective cylinder, such that their transfer resistance is identical in both models. In the reduced model, these synapses are merged into one “NEURON” process (red synapse in **Fig. 1b**). However, they retain their individual activation times (see **Methods**). **Figure 1d** compares the transfer impedance between a specific point in the apical tree (marked by ‘**d**’ in **Fig. 1a and b**) and the soma. By construction, for the passive case, the transfer resistance (for ω = 0) is equivalent for the respective loci in the detailed and the reduced model. This is indeed the case in **Fig. 1d** (left-most point on the x-axis), thus validating the implementation of the *Neuron_Reduce* analytic method. Note that although constructed using ω = 0, the similarity between the detailed and reduced model also holds for higher input frequencies. However, for *ω* around 10 - 100 Hz, the transfer impedance from d-to soma (and vice versa, due to the reciprocity theorem for passive systems^43^) is somewhat larger in the reduced model (compare the red and black lines).

To test the performance of *Neuron_Reduce* on transient synaptic inputs (composed of mixed input frequencies), we sequentially activated the four synapses shown in **Fig. 1c** in both the detailed and the reduced models (see **Methods** and Supplementary Table 2). **Figure 1e** shows the close similarity in the composite somatic EPSPs between the two models, further validating that the mapping of the detailed model to the reduced model using ω = 0 provides satisfactory results for the passive case (see also Supplementary Fig. 2).

**Figure 2.**
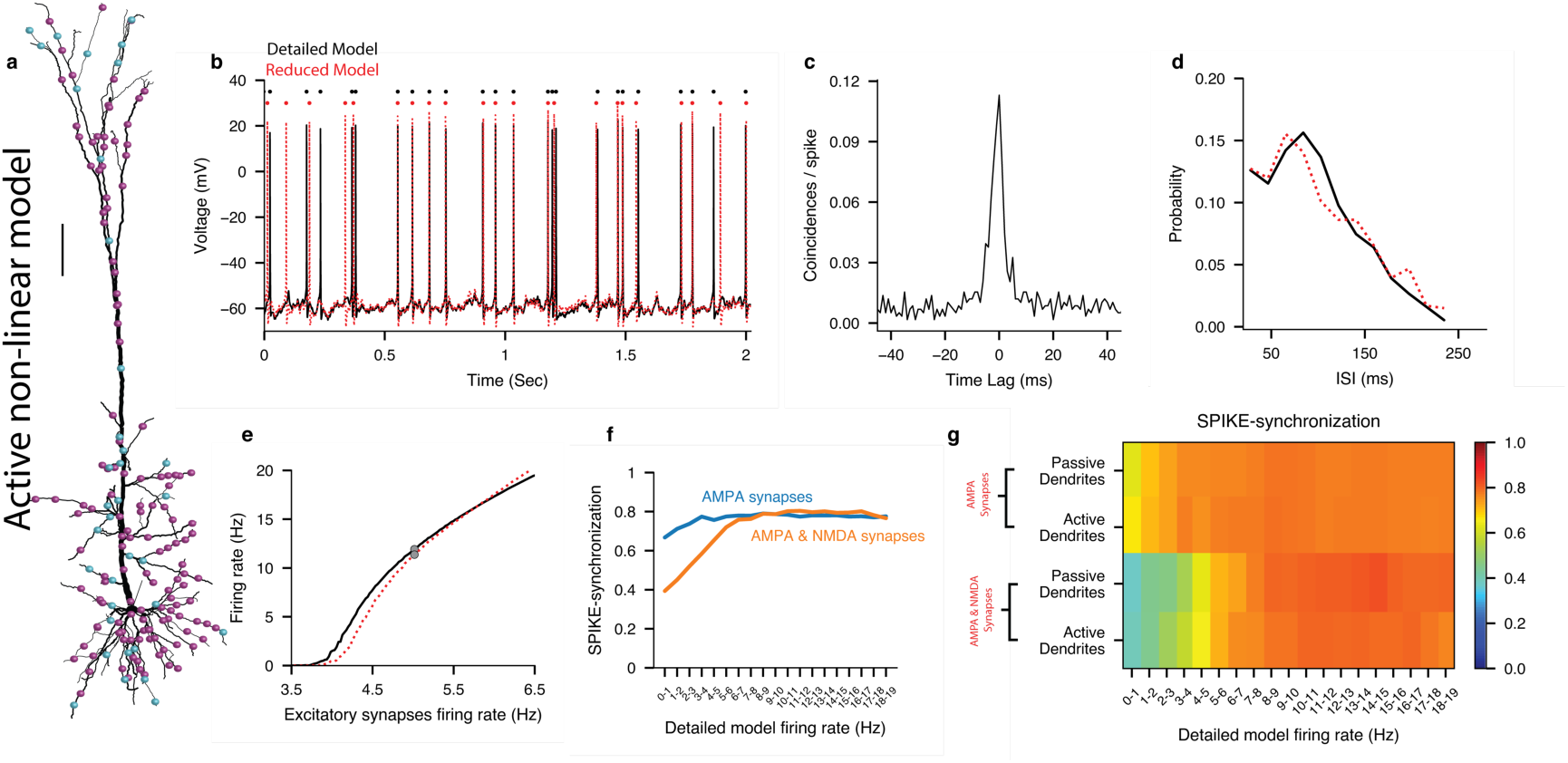
*Neuron_Reduce* faithfully replicated the I/O properties of a detailed nonlinear model of a L5 pyramidal cell. **a** Layer 5 pyramidal cell model^5^ as in Fig. 1a, with 8000 (AMPA + NMDA) excitatory (magenta dots) and 2000 inhibitory synapses (cyan dots, see Supplementary Table 2 for synaptic parameters). Excitatory synapses were activated randomly at 5Hz and the inhibitory synapses at 10 Hz. This detailed model consists of a dendritic *Ca^2+^* ^“^hot spot” (as in Fig. 1c) and an *Na^+^* spiking mechanism at the cell body. Scale bar 100 µm. **b** An example of the voltage dynamics at the soma of the detailed model (black trace) and the reduced model (red trace); spike times are represented by the black and red dots above the respective spikes. **c** Cross-correlation between spikes in the reduced versus the detailed models. **d** Inter-Spike Interval (ISI) distributions for the two models. **e** Output firing rate of the reduced (red) versus the detailed (black) models as a function of the firing rate of the excitatory synapses. Grey dots represent the case shown in b. **f** SPIKE-synchronization measure between the two models as a function of the firing rate of the detailed model for the case of only AMPA (blue) and AMPA + NMDA synapses (orange). The performance of the reduced model with NMDA-synapses was lower for low output frequency but improved significantly for output frequencies above ∼7 Hz (see **Discussion**). **g** SPIKE-synchronization between the detailed and the reduced models as a function of the firing rate of the detailed model, for active and passive dendrites, and with/without NMDA-based synaptic conductance.

### Accuracy and speed-up of *Neuron_Reduce* for nonlinear neuron models

To measure the accuracy of *Neuron_Reduce* for a fully-active nonlinear neuron model we ran a comprehensive set of simulations using the well-established case of the L5 pyramidal cell model^5^ shown in **Fig. 2a** (same cell as in **Fig. 1)**. This neuron model includes a variety of nonlinear dendritic channels including a voltage-dependent *Ca^2+^* “hot spot” in the apical tuft (schematic yellow region in **Fig. 1c**) and an *Na^+^*-based spiking mechanism in the cell body. We randomly distributed 8,000 excitatory and 2,000 inhibitory synapses on the modeled dendritic tree (the synaptic parameters are listed in Supplementary Table 2) and used *Neuron_Reduce* to generate a reduced model for this cell. We simulated the detailed model by randomly activating the excitatory synapses at 5 Hz and the inhibitory synapses at 10 Hz (see **Methods**). The detailed model responded with an average firing rate of 11.8 Hz (black trace in **Fig. 2b**; only 2 sec out of 50 sec simulation time are shown). The average firing rate of the respective reduced model in response to the same synaptic input was 11.3 Hz (red trace, **Fig. 2b**; spike timings are shown by small dots on the top). The cross-correlation between the two spike trains peaked around zero (**Fig. 2c**), and the inter-spike interval distributions of the two models were similar (**Fig. 2d**).

The full range of responses to a random synaptic input for the two models was explored by varying the firing rate of the excitatory (AMPA- and NMDA-based) synapses and measuring the degree of similarity between the firing rates of the two models, which indicated a good fit between the two (**Fig. 2e**). We used the *SPIKE-synchronization* measure^44, 45^ to further quantify the similarity between the spike trains of the detailed and reduced models. The SPIKE-synchronization value for the two spike trains shown in **Fig. 2b** was 0.8. In **Fig. 2f**, the SPIKE-synchronization was computed as a function of the output rate of the detailed model for both the case where the excitatory synapses consisted of only an AMPA-component (blue) and for when they also consisted of an NMDA component (orange). For the AMPA-only case, the SPIKE-synchronization was high for all output frequencies but was low for low output frequencies when the synapses consisted of an NMDA component, although improving significantly for output frequencies above ∼7 Hz (see **Discussion**). **Figure 2g** shows the SPIKE-synchronization as a function of the firing rate of the detailed model, for active and passive dendrites and with/without NMDA-based synaptic conductance, demonstrating again that when NMDA synapses are involved, the performance of the reduced model is low for low output rates. We also tested other spike trains similarity metrics^46, 47^ (Supplementary Fig. 3) and found comparable results to those shown in **Fig. 2**. We have also analyzed the performance of *Neuron_Reduce* on two additional patterns of synaptic input. In one case, the synaptic input was activated in an oscillatory manner at different frequencies (see **Methods**). In these cases, the spike synchronization measure ranged between 0.75 - 1 (**Supplementary Fig. 4a, b**). In the other case the synaptic input was taken from a spontaneously active Blue Brain circuit^17^ (see **Methods**). In this case the spike synchronization measure was 0.708 (Supplementary Fig. 4c).

**Figure 3.**
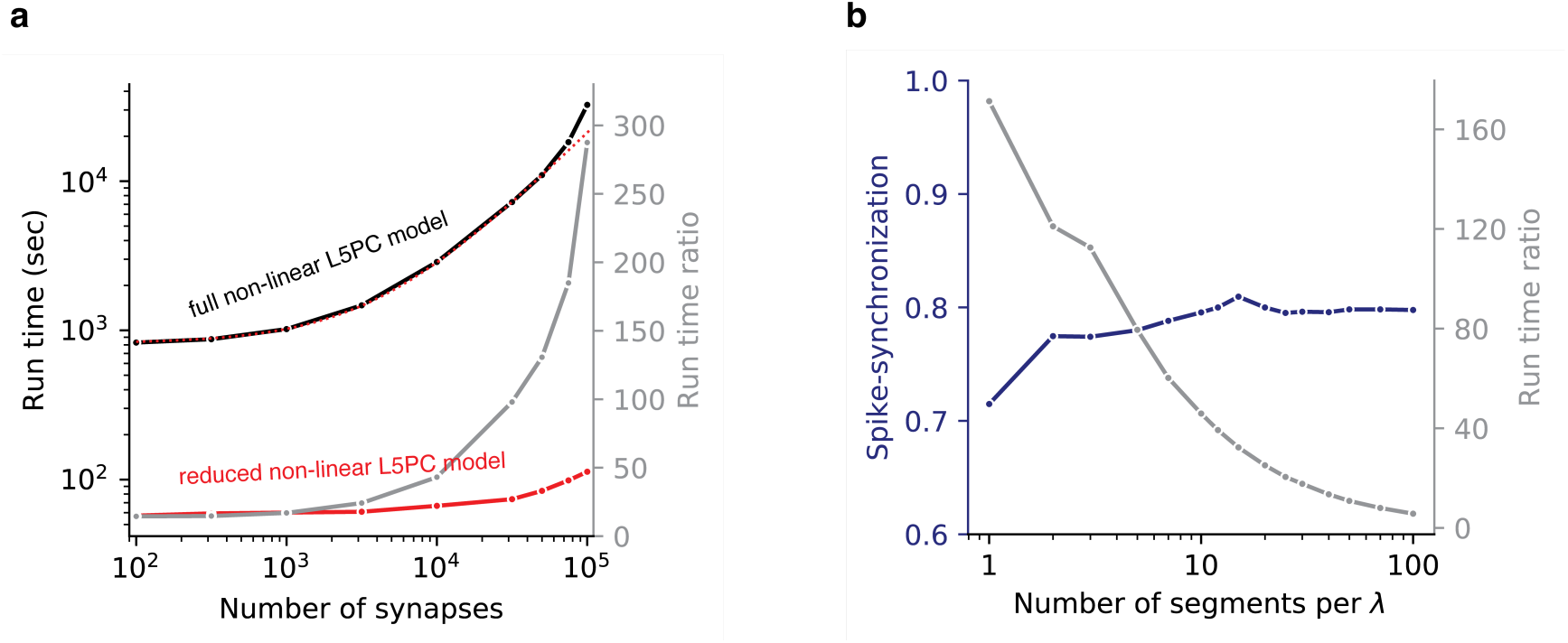
*Neuron_Reduce* enhances the simulation speed by up to hundreds-fold. **a** Simulation run-time for the detailed (black) and the reduced models (red) of layer 5 pyramidal cell shown in Fig. 2a, for a simulation of 50 seconds, and their ratio (the speed-up, grey) as a function of the number of simulated (GABAA-, AMPA- and NMDA-based) synapses. Due to the almost constant run-time of the reduced model, the run-time ratio increases with larger number of synapses. Above 75,000 synapses, an additional effect becomes visible: the detailed model no longer fits into the cache of the CPU and exhibits a supralinear increase in runtime. This can be seen by the black curve deviating from the dotted red curve, which shows the expected simulation time for the detailed model assuming a constant computation cost per synapse (see also Supplementary Table 1). **b** Accuracy (blue) of the reduced model and its speed-up in simulation run-time (grey) as a function of the number of electrical compartments per length constant for a neuron with 10,000 synapses (50 seconds per simulation)

**Figure 4.**
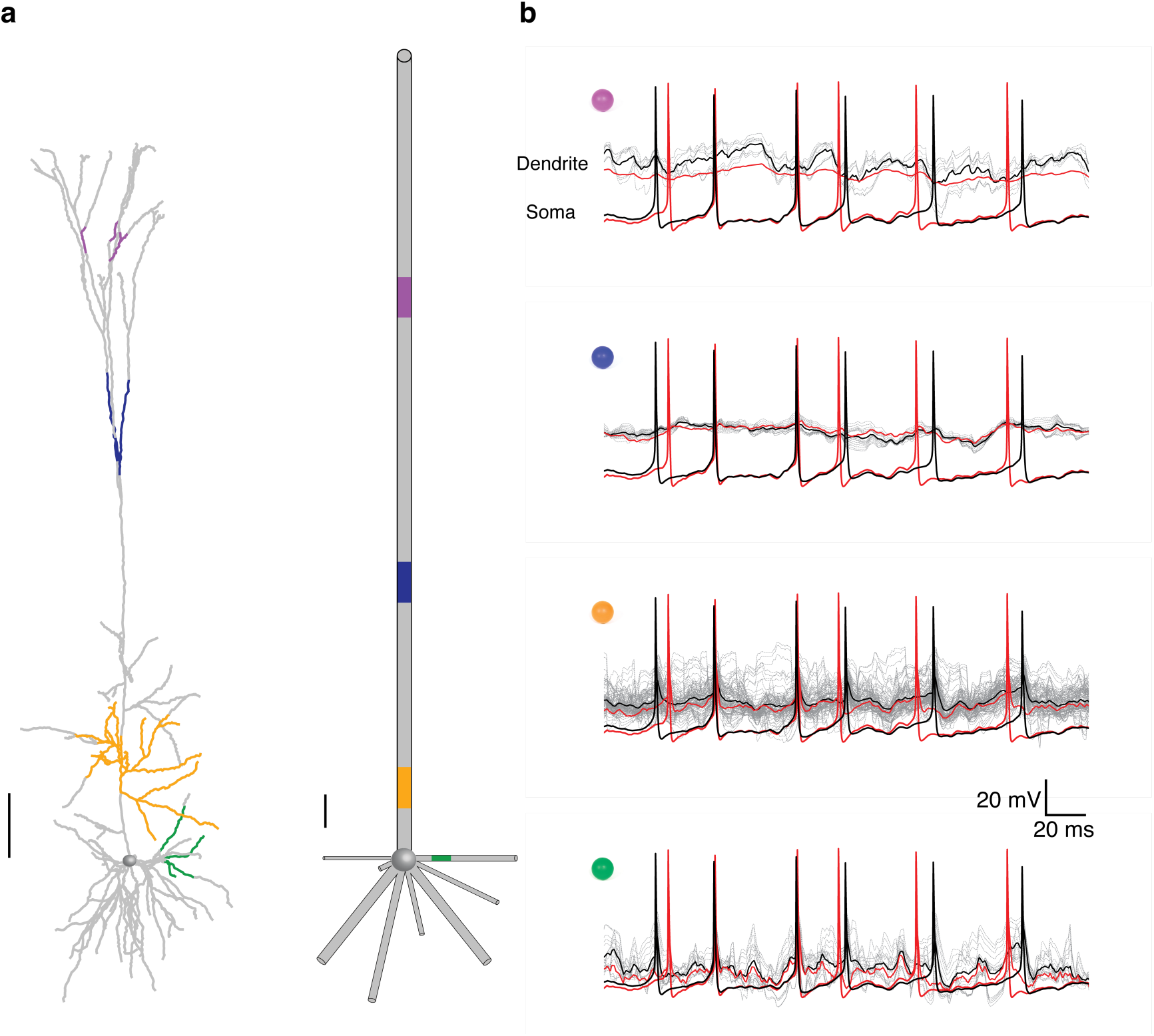
The dendritic potential in the reduced model represents the average dendritic voltage dynamics in the detailed model. **a** Detailed model (left) and reduced model (right) of the cell shown in Fig. 2. Dendritic branches of the same color in the detailed model are all mapped to the respective compartment with identical color in the reduced model. **b** For each of the four colored regions shown in a (and respective colored sphere at top left), the voltage transients in individual branches are shown by the grey traces. Superimposed in black is the average voltage of these traces and in red is the voltage transient in the respective compartment in the reduced model. The somatic spikes in the detailed model (black) and reduced model (red) are also shown. The simulation is as in Figure 2e, with excitatory synapses firing at 5.5 Hz. Scale bars for the respective morphologies are 100 µm.

We compared the performance of our reduction method to two other reduction approaches, one of which was Rall’s “equivalent cable” reduction method^31, 48^. The other method maps all the dendritic synapses to the somatic compartment, after computing the filtering effect of the dendritic cable for each synapse^34^ (see **Methods**). *Neuron_Reduce* outperformed both these reduction methods (Supplementary Fig. 5).

**Figure 5.**
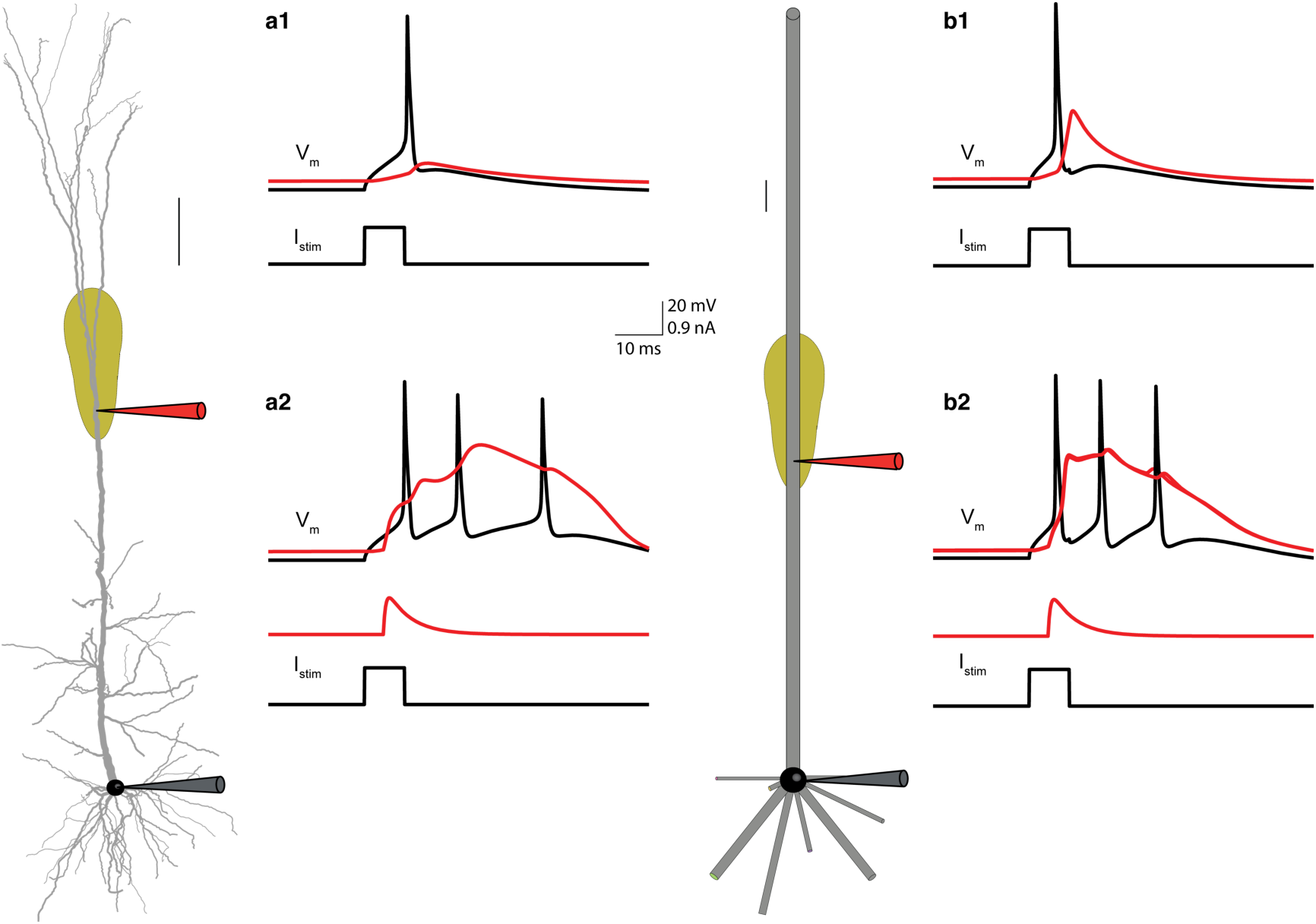
Dendritic Ca^2+^-spike and BAC-firing faithfully replicated in the reduced model. a1-a2. Left, detailed L5 pyramidal cell model with nonlinear *Ca^2+^* ^“^hot spot” (same model as in Fig. 2). **a1** Injecting a depolarizing step current to the soma (0.95 nA for 8.5 ms) in the detailed model evoked a somatic action potential, AP (black trace) that propagated backward semi-actively into the apical tree (red trace). **a2** Combining the somatic input with a transient synaptic-like current injection (0.95 nA peak value with 0.5 ms and 5 ms rise-time and decay-time respectively) to the “hot region” in the apical dendrite evoked a prolonged local *Ca^2+^* spike which, in turn, triggered a burst of two extra somatic *Na^+^* spikes (the BAC firing phenomenon^50^). **b1-b2** Same as in a1-a2, but for the reduced model. Scale bars for the respective morphologies are 100 µm.

**Figure 3** compares the run-time of the detailed versus the reduced model for the neuron model shown in **Fig. 2a**. Simulating the detailed model with 10,000 synapses for 50 seconds of biological time required 2,906 seconds of computer time (run-time), whereas it took only 68.7 seconds in the reduced model, a ∼42-fold computational speed-up (see Supplementary Table 1). The larger the number of synapses in the detailed model, the longer the run-time (**Fig. 3a**). In contrast, the run-time in the reduced model is only shallowly dependent on the number of synapses. This is expected when considering the synaptic merging step in our algorithm (see **Discussion**). The run-time of the reduced model depends on the number of compartments per cylinder; it increases sharply with an increasing number of compartments (the run-time ratio between the detailed and the reduced models decreases, grey line in **Fig. 3b**). However, there was no improvement in the SPIKE-synchronization measure when the spatial discretization, ΔX, per compartment was less than 0.1λ (**Figure 3b** blue line and see also previous research on the subject^49^). Therefore, all the results presented in **Fig. 1-7** are based on models with a ΔX that does not exceed 0.1λ.

In **Figure 4** we compared the dendritic voltage in the detailed and in the respective location in the reduced model. We found that: (i) the voltage transients could differ significantly in dendritic branches that are all mapped to the same compartment in the reduced model (e.g., compare the grey traces in the yellow compartments in **Fig. 4b**). (ii) the average voltage trace of these different dendritic branches (black trace in **Fig. 4b**) is similar to the voltage in the respective compartment in the reduced model (red trace in **Fig. 4b**). The implications of the latter finding for capturing highly nonlinear local dendritic events is elaborated in the **Discussion**.

### *Neuron_Reduce* replicates key dendritic nonlinearities and computations

To determine the capabilities of the reduced models to support nonlinear dendritic phenomena and dendritic computations, we repeated two classical experiments in both the detailed and the reduced model of the L5 pyramidal cell shown in **Fig. 1**. The first simulated experiment started by injecting a brief depolarizing step current to the soma of the detailed model to generate a somatic *Na^+^* action potential (AP, black trace in **Fig. 5a**). This AP propagated backward to the apical dendrite, the BPAP (red trace in **Fig. 5a1**). Repeating the same current injection in the reduced model led to a similar phenomenon, but with a larger BPAP (**Fig. 5b1**). The detailed model also included a “hot region” with voltage-dependent calcium conductances in its apical dendrite (see also **Fig. 1**). Combining somatic current injection with synaptic-like transient depolarizing current injected to the apical nexus evoked a prolonged *Ca^2+^* spike in the distal apical dendrite (red trace at the apical tree) which, in turn, generated a burst of somatic *Na^+^* spikes (the BAC firing^4, 5, 50^, **Fig. 5a2**). *Neuron_Reduce* maps the nonlinear dendritic “hot” *Ca^2+^* region to its respective location in the reduced model (see **Fig. 1** and **Methods**). **Fig. 5b1-b2** shows that the exact same combination of somatic and dendritic input currents also produced the BAC firing phenomenon in the reduced model. However, the reduced model was somewhat more excitable than the detailed model; this resulted in a burst of three spikes with a higher frequency (and sometimes with an additional spike) in the reduced model (compare **Fig. 5a2 to b2**).

The second simulated experiment attempted to replicate theoretical and experimental results reported in previous studies^1, 51, 52^. In these studies, several excitatory synapses were activated sequentially in time, on a stretch of a basal dendrite, either in the soma-to-dendrites (OUT) direction or vice versa (the IN direction). Rall showed that the shape and size of the resultant composite somatic EPSP depended strongly on the spatio-temporal order of synaptic activation; it was always larger and more delayed for the centripetal (dendrites-to-soma) than for the centrifugal (soma-to-dendrites) sequence of synaptic activation (this difference can serve to compute the direction of motion^51^). It was shown that the difference in the resulting somatic voltage peak between these two spatio-temporal sequences of synaptic activation was enhanced when nonlinear NMDA-dependent synapses were involved and that it made it possible to discriminate between complex patterns of dendritic activation^52^.

To simulate these phenomena, twelve excitatory synapses were placed along one basal branch in the detailed model (red dots on the green basal tree, **Fig. 6a1**). At first, the synapses only had an AMPA component. The synapses were activated in temporal order from the tip to the soma (IN, cyan traces) or from the soma to the tip (OUT, blue traces, see **Methods** for details). As predicted by Rall, activation in the IN direction resulted in larger and delayed somatic EPSP (cyan trace versus the blue trace in **Fig. 6a2**). *Neuron_Reduce* merged these twelve synapses into five point processes along the respective cylinder (**Fig. 6b1**). We repeated the same experiment in the reduced model and found that the EPSP resulting from the IN direction was larger and delayed, with a similar EPSP waveform to that of the detailed model (see also Supplementary Fig. 2 and **Discussion**). Next, an NMDA component was added to the 12 simulated synapses; this resulted in larger somatic EPSP amplitudes in both directions (and both models) and a smaller difference in the peak timing between the different directions in both the detailed and the reduced model (compare **Fig. 6a3 to b3**).

To generalize the impact of the spatio-temporal order of synaptic activation, we used a directionality index suggested in a previous study^52^. This measure estimates how different a given synaptic sequence is from the IN sequence by calculating the number of synaptic swaps needed to convert this given pattern into the IN pattern (using the bubble-sort algorithm, see **Methods**). We tested the EPSPs that resulted from different temporal combinations of synaptic activation (each having a different directionality index), both without (**Fig. 6c1**) and with an NMDA component (**Fig. 6d1**). The peak somatic EPSP in the reduced model (red dots) was larger than in the respective detailed model (black dots), both for the AMPA-only case (by 1.71 ± 0.43 mV) and for the AMPA + NMDA case (by 4.80 ± 0.74 mV); see **Supplementary Fig. 1**. Nevertheless, the behavior of the two models was similar when the somatic voltage in the two models was subtracted by the peak value obtained in the OUT direction (**Fig. 6c2 and 6d2**). Then, the difference between the reduced and the detailed models was, on average, only 0.11 ± 0.43 mV for the AMPA-only case and 0.35 ± 0.43 mV for the AMPA + NMDA case. Thus, although the detailed and the reduced models differ to a certain extent (see **Discussion**), the capability of the reduced model to discriminate between spatio-temporal patterns of synaptic activation is similar to that of the detailed model.

### *Neuron-Reduce* can be implemented successfully on a variety of neuron models

We next tested the utility of *Neuron_Reduce* on 13 different neuron models from different brain regions (**Fig. 7**). Four models were obtained from the Blue Brain database^17, 53^: L6 Tufted Pyramidal Cell, L4 Double Bouquet Cell, L4 Spiny Stellate Cell and L5 Martinotti cell from the rat somatosensory cortex. Two additional models were obtained from the Allen Institute cell type database^11^: an L4 Spiny cell and an L1 Aspiny Cell from the mouse visual cortex. Medium spiny neuron from the mouse basal ganglia^54^; two rat thalamocortical neurons^55^; Golgi cell from mouse cerebellar cortex and one inhibitory hippocampal neuron from the rat ^56^. We also took two additional neuron models from our laboratory: rat L2/3 large basket cell^57^ and a model of a human L2/3 pyramidal cell from the temporal cortex^58^. All these models were based on 3D reconstructions and were constrained by experimental recordings (see Supplementary Table 2 for details on the various neuron models and input parameters).

*Neuron_Reduce* successfully generated a reduced model for all these different cell types, with highly faithful response properties in all cases (**Fig. 7**). Three examples with their respective morphologies for the detailed and reduced models are shown in **Fig. 7a-c**. For a given input we measured the spiking activity of the detailed and reduced models (**Fig. 7d-f**) and calculated the corresponding SPIKE-synchronization values. For the L6 tufted PC model (**Fig. 7a, d**), the L2/3 large basket cell model (**Fig. 7b, e**) and the L6 double bouquet model (**Fig. 7c, f**) the SPIKE-synchronization values were 0.74, 0.85, and 0.91 respectively, for 50 second long simulations (only 2 seconds are shown in **Fig. 7d-f**). The SPIKE-synchronization values for additional inputs, and for the other 10 neuron models and their corresponding reduced models, are shown in **Fig. 7g**. We have also tested the performance of *Neuron_Reduce* and the variability of the SPIKE-synchronization measure using eight neocortical neuron types, with 11 cell models per type taken from the Blue Brain cells dataset^17, 53^. Supplementary Figure 6 shows that, for all cells the SPIKE-synchronization measure remains similar to that found in **Figure 7**, mean values per cell type ranging between 0.43-0.86, increasing with output frequency of the modelled cell.

**Figure 6.**
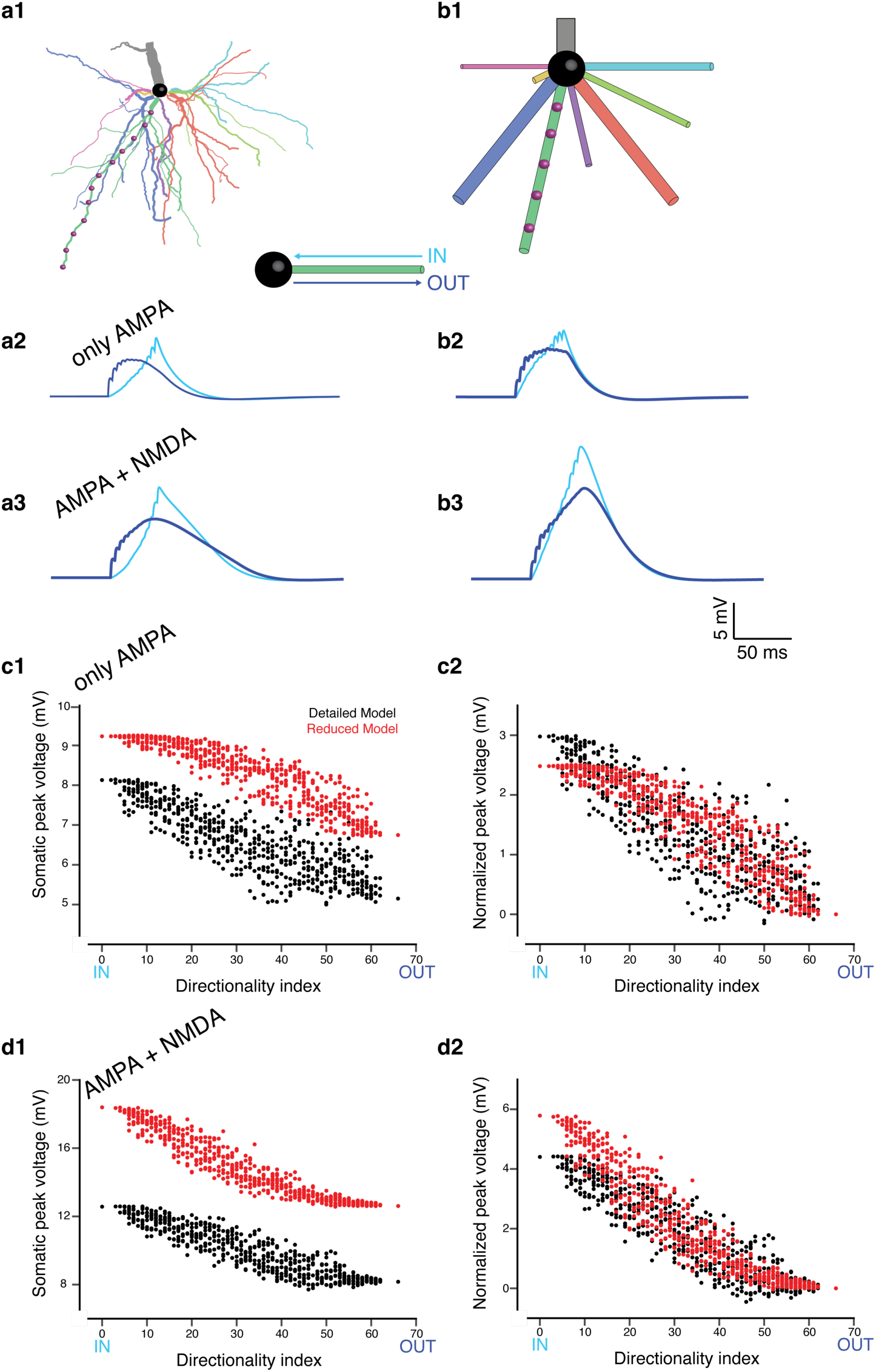
Discriminating spatiotemporal input sequences in the detailed versus the reduced model. a1. A model of L5PC (detailed model, Fig. 1) with twelve excitatory synapses spatially distributed on one of its basal dendrites (red dots on green basal dendrite). **a2** Somatic responses to sequential activations of its basal synapses in the IN (cyan) and the OUT (blue) directions. In this case, the synaptic model only consists of an AMPA component. **a3** As in a2 but the synaptic model consists of both AMPA and NMDA components. **b1** Reduced model for the detailed model shown in a1. *Neuron_Reduce* mapped the twelve synapses in the detailed model into five synapses in the reduced model. **b2 and b3**. As in a2 and a3, but for the reduced model. **c1** Pattern separability (see **Methods**) of the detailed (black) and the reduced (red) models when the synaptic model only consists of an AMPA component. **c2** As in **c1**, after subtracting the peak voltage obtained in the OUT direction from each of the voltage responses. **d1 and d2** As in c1 and c2 but when the synaptic models consisted of both AMPA- and NMDA-conductances. Note the similarity between the detailed and the reduced model in terms of pattern separability.

**Figure 7.**
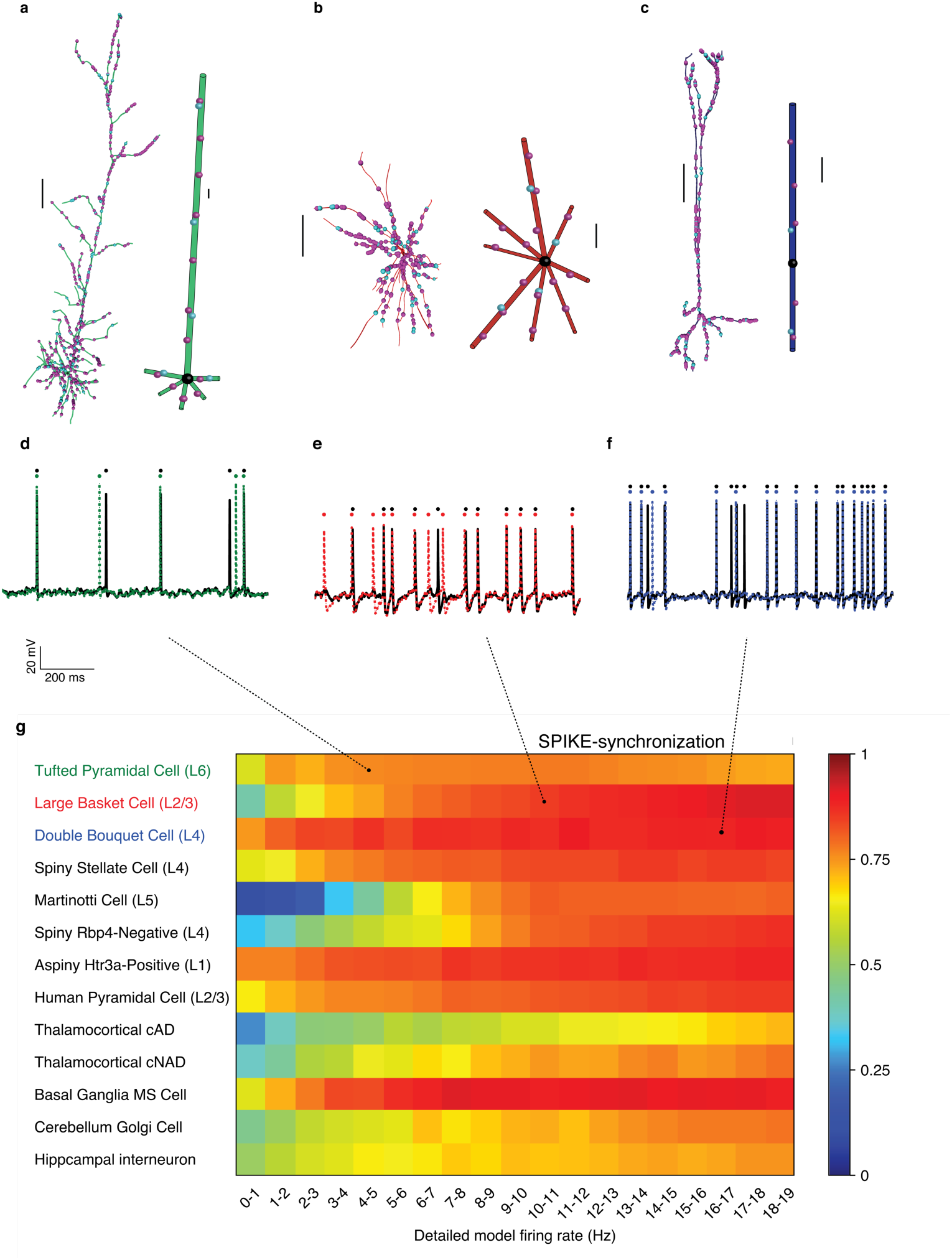
*Neuron_Reduce* working successfully on a variety of neuron models. a-c. Detailed models of three somatosensory neurons (left, L6 Tufted Pyramidal Cell in green; middle, L2/3 Large Basket Cell in red; and right, L4 Double Bouquet Cell in blue) and their respective reduced models. Scale bars 100 µm. **d-f** Voltage responses to an excitatory synaptic input activated at 1.8, 2.9, 3.17 Hz respectively for both the detailed (black) and the reduced models (corresponding colors). The inhibitory input activation rate was 10 Hz for all models. **g** The SPIKE-synchronization index for the 13 detailed versus reduced neuron models. The mean simulation speed-up for the L6 Tufted Pyramidal cell, L5 Martinotti cell and L4 Spiny Stellate cell were 95, 40 and 60, respectively. See Supplementary Table 2 for cell models and input parameters and Supplementary Figure 6 for the SPIKE-synchronization measure on additional 88 modeled cells.

## Discussion

### *Neuron_Reduce*: theoretical underpinning and advantages

*Neuron_Reduce* is a new tool for simplifying complex neuron models while enhancing their simulation run-time. It analytically maps the detailed tree into a reduced multi-cylindrical tree, based on Rall’s cable theory and linear circuit theory (**Fig. 1**). The underpinning of the reduction algorithm is that it preserves the magnitude of the transfer impedance |*Z*_0,*j*_(*ω*)| from each dendritic location, j, to the soma (the dendro-somatic direction, Eqs. (1) – (11) in **Methods**). Since in linear systems, it holds that |*Z*_0,*j*_(*ω*)| = |*Z_j_*_,0_(*ω*)|, for passive dendritic trees it also preserves the transfer impedance in the somato-dendritic direction (e.g., current injection at the soma will result in the same voltage response at the respective sites in the detailed and reduced models^59^).

Note that dendritic voltage transients (e.g., synaptic potentials) contain a range of frequencies, *ω*. We however had to select one frequency to use for the mapping of the detailed-to-the-reduced tree. Consequently, we examined a whole range of possible *ω* values for this mapping. Conveniently we found that ω = 0 is the preferred frequency for generating the reduced model (namely, when the mapping from detailed to reduced model is performed based on the transfer resistance |*Z*_0,*j*_(*ω*=0)| = |*R*_0,*j*_|) see Supplementary Fig. 7. This result is actually not surprising; Rinzel and Rall^33^ showed that, in passive trees and current-based synapses, the attenuation of the voltage time-integral (the area below the EPSPs) is identical to the attenuation of steady-state voltage. In other words, when using the transfer resistance for our mapping procedure, we preserved the total charge transfer (which in our case, was proportional to the voltage time integral) from the synapse to the soma (and vice versa), but not e.g., the EPSP peak value.

*Neuron_Reduce* was proven to be accurate in replicating voltage dynamics and spike timing for a large regime of input parameters and a variety of neuron types (**Fig. 7**, Supplementary Figure 6 **and** Supplementary Table 2). This claim is based on using several metrics for assessing the quality of the performance of the reduced model (Supplementary Fig. 3). Moreover, we implemented STDP rule on synaptic weights in both the detailed and reduced model and found great similarity in the weight trajectories of the respective synapses (not shown). *Neuron_Reduce* is straightforward to use, it is fast and generally applicable, thus enabling its implementation on any neuron morphology with any number (even tens of thousands) of synapses. One key advantage of *Neuron_Reduce* is that it retains the identity of individual dendrites and synapses and that it maps dendritic nonlinearities to their respective loci in the reduced model, hence preserving local excitable dendritic phenomena and therefore maintaining nonlinear dendritic computations. *Neuron_Reduce* also preserves the passive cable properties (*Rm, Ra, and Cm*) of the detailed model, thus preserving synaptic integration and other temporal aspects of the detailed model. We have also examined *Neuron_Reduce* for the case of a gap junction connecting dendrites of two L2/3 large basket cells. As *Neuron_Reduce* preserves the transfer resistance from the location of the synapses (in this case the gap junction) to the soma and *vice versa,* one expects that the coupling coefficient between the two connected cells will be preserved in the reduced models, after mapping the gap junction to its appropriate location in the reduced model. This is indeed what we have found (not shown).

*Neuron_Reduce* enhances the computational speed by a factor of up to several hundred folds, depending on the simulated morphology and the number of simulated synapses (**Fig. 3** and Supplementary Table 1). This combination of capabilities, together with its user-friendly documentation and its public availability, make *Neuron_Reduce* a promising method for the community of neuronal modelers and computational neuroscientists, and for the growing community interested in “biophysical deep learning”.

For a large number of synapses and complex morphologies, the run-time of *Neuron_Reduce* models can be accelerated by up to 250 fold as compared to their respective detailed models (**Fig. 3** and Supplementary Table 1). This is achieved in two associated steps. First, the algorithm reduces the number of compartments of the neuron model; e.g., for the reconstructed tree in **Fig. 1**, it reduced the number of compartments from 642 to 50. Then, synapses (and ion channels) that are mapped to the same electrical compartment in the reduced tree (because they have similar transfer resistance to the soma) are merged into one point process in NEURON. Each of these steps on its own has a relatively small effect on the run-time. However, when combined, a large (supra-linear) improvement in the computational speed is achieved (Supplementary Table 1). This is because at each time step, NEURON computes both the voltage in each electrical compartment as well as the currents and states of each point process and membrane mechanism (synapses and conductances). Reducing the number of compartments in a model decreases the number of equations to be solved and the number of synapses to be simulated (due to the reduced number of compartments, a larger number of synapses are merged together). Importantly, merging synapses preserves the activation time of each synapse. Note, however, that in its present state, *Neuron_Reduce* cannot merge synapses with different kinetics.

### Other reduction methods

Several other reduction methods for single neurons have been proposed over the years^12, 34–39^. Most are not based on an analytic underpinning and thus require hand-tuning of the respective biophysical and morphological parameters. In addition, most of these methods have not been examined using realistic numbers of dendritic synapses and are incapable of systematic incorporation of dendritic nonlinearities. In most cases, their accuracy has not been assessed for a range of neuron types (but see ^41^). Many of these methods are not well-documented, thus making it hard to compare them directly with *Neuron_Reduce*. Nevertheless, we did compare the performance of *Neuron_Reduce* to two other reduction methods and showed that it outperformed them (Supplementary Fig. 5).

### Limitations of *Neuron_Reduce* and future work

It should be noted that although the transfer impedance from a given dendritic locus to the soma is preserved in the reduced model, the input impedance at that locus is not preserved (is lower) in the reduced model. Consequently, the conditions for evoking local dendritic events, and the fine details of these events are not identical in the detailed and the reduced models (e.g., compare **Fig. 5a1 and a2** to **Fig. 5b1 and b2 and see Fig. 4**). Indeed, if there were highly local dendritic Na^+^ spikes (as in^60^) then *Neuron_Reduce* will not capture them, as this local dendritic spike will be averaged-out in the respective lumped cable. Similarly, because the local voltage response to a current injection in the dendrite depends on the dendritic impedance, the local synaptic responses are somewhat different in the detailed versus the reduced cases, especially when voltage-gated ion channels (such as NMDA-dependent synaptic channels) are involved. In fact, when large dendritic NMDA signals are involved, the resultant somatic EPSPs are expected to be different in the detailed as compared to the reduced model, as is the case in **Figures 2** and **6**. Indeed, if one insists on preserving highly local nonlinear dendritic events, then the full dendritic tree should be modeled.

Despite these local differences, the reduced model for L5PC did generate a local dendritic *Ca^2+^* spike in the cylinder representing the apical dendrite and was able to perform an input classification task (enhanced by NMDA-conductance), as in the detailed tree (**Figures 5** and **6**). Moreover, when embedded in large circuits, individual neurons are likely to receive semi-random dendritic input, rather than a clustered input on specific dendrites. For such inputs, the reduced models generated by *Neuron_Reduce* capture most of the statistics of the membrane voltage dynamics as in the detailed model (**Figures 2** and **7 and** Supplementary Figures 6 and 4).

The next straightforward step is to use *Neuron_Reduce* to simplify all the neurons composing a large neural network model, such as in the Blue Brain Project^17^, the *in silico* model presented by Egger et al^16^ and by Yazan et al^61^. By preserving the connectivity and reducing the complexity of the neuronal models, the reduced models will make it possible to run much longer simulations and/or larger neuronal networks, while faithfully preserving the I/O of each neuron. Such long simulations are critical for reproducing long-term processes such as circuit evolution and structural and functional plasticity.

## Methods

### *Neuron_Reduce* algorithm and its implementation in NEURON

*Neuron_Reduce* maps each original stem dendrite to a unique single cylinder with both ends sealed. This cylinder preserves the specific passive cable properties (*R_m_, C_m_*, and *R_a_*) of the original tree as well as both the transfer impedance from the electrotonically most distal dendritic tip to the soma and the input resistance at the soma end of the corresponding stem dendrite (when disconnected from the soma). For a sinusoidal angular frequency ω > 0, the transfer impedance *Z_i,j_(ω)* is the ratio between the Fourier transform of the voltage at point (*i*) and the Fourier transform of the sinusoidal current injected into the injection point (*j*) (note that in passive systems, *Z_i,j_(ω)* = *Z_j,i_(ω)*). This ratio is a complex number; its magnitude (|*Z_i,j_(ω)*|) is the ratio (in Ω) between the peak voltage response and the amplitude of the injected current. In a short cylindrical cable with sealed ends and electrotonic length *L*, the transfer impedance, *Z_0,X_(ω)*, between the somatic end of the cylinder (X = 0) and any location *X* is^62, 63^

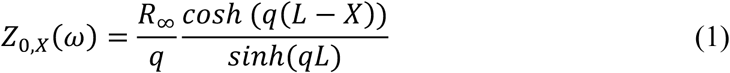

where

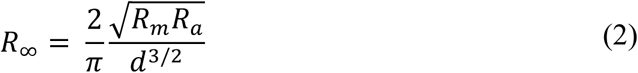

and

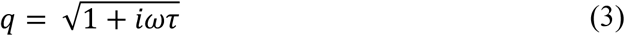

where *τ* is the membrane time constant, *R_m_C_m_*.

From Eq. (1), the input impedance at *X* = 0 is

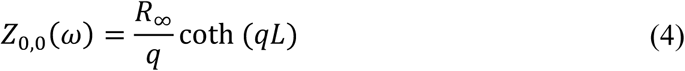

We next want a cylindrical cable of electrotonic length *L*, in which both |*Z*_0,*L*_(*ω*)| and |*Z*_0,0_(*ω*)| are identical to those measured in the respective original stem dendrite (**Fig. 1**). For this purpose, we first look for an *L* value in which the ratio |*Z*_0,*L*_(*ω*)|/|*Z*_0,0_(*ω*)| is preserved. Dividing Eq. (1) by Eq. (4) we get

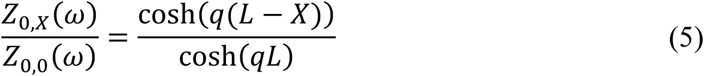

which can be expressed as

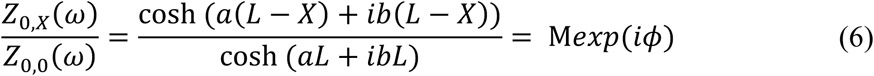

where *a* and *b* are the real and the imaginary parts of *q*, respectively, and M and *ϕ* are the modulus and phase angle of this complex ratio.

As shown previously^62^ it follows that

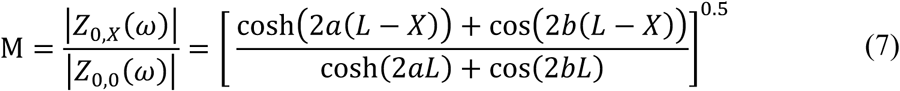

and

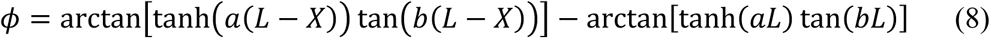

Importantly, for a fixed M (and a given *ω* there is a unique value of *L* that satisfies Eq. (7) (see Figure 4 in ^62^ and note the-one-to-one mapping between M and L for a given *ω* value). However, there are an infinite number of cylindrical cables (with different diameters and lengths) that have identical *L* values preserving a given M value in Eq. (7).

We next need a unique cable, with a specific diameter *d*, that also preserves the measured value of |*Z*_0,0_(*ω*)| (and therefore it also preserves |*Z*_0,*L*_(*ω*)|, see Eq. (7)).

From Eqs. (2) and (4) we get

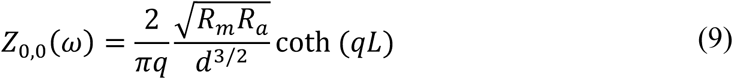

and thus

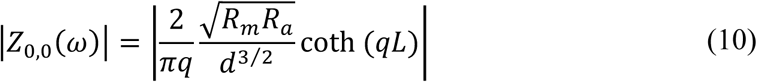

from which we compute the diameter, *d*, for that cylinder

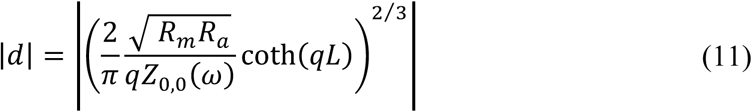

Equations (1) – (11) provide the unique cylindrical cable (with a specific *d* and *L*, and the given membrane and axial properties) that preserves the values of |*Z*_0,*L*_(*ω*)| and |*Z*_0,0_(*ω*)| as in the respective stem dendrite. Note that this unique cable does not necessarily preserve the phase ratio (*ϕ* in Eq. (8)) as in the original tree.

Practically, in order to transform each original stem dendrite (with fixed *Rm, Ra, and Cm* values) into a corresponding unique cylindrical cable, we proceeded as follows. First, on each modeled stem dendrite (when isolated from the soma), we searched for a distal location *x* with minimal transfer impedance, |*Z*_0,*x*_(*ω*)|, from that particular *x* to the soma. This location provided the smallest M value for this particular stem dendrite. This distal dendritic locus, *x,* was mapped to the distal end, X = L, of the corresponding cylinder. We then used Eqs. (1) – (11) to calculate the unique respective cylinder for each stem dendrite.

In order to map synapses from the detailed model to the reduced one we computed, for each synapse at location *j* in the detailed model, |*Z*_0,*j*_(*ω*)|, and then mapped this synapse to the respective location *y* in the reduced model, such that |*Z*_0,*y*_(*ω*)| = |*Z*_0,*j*_(*ω*)|. This reduced model is then compartmentalized into segments (typically with spatial resolution of 0.1λ, see **Fig. 3b**). We then merged all synapses with identical kinetics and reversal potential, that are mapped to a particular segment, onto a single point process object in NEURON (**Fig. 1**, step B). These synapses retain their original activation time and biophysical properties through the connection of each of their respective original NetStim objects to the single point process that represents them all (each of these connections was mediated by the synapse’s original NetCon object). As shown in Supplementary Table 1, this step dramatically reduced the running time of the model. We note that all the results presented in this study were obtained using *ω* = 0 in Eqs. (1) - (11), since running the same simulations with *ω* = 0 provided the best performance (see Supplementary Fig. 7). However, *ω* is a parameter in the algorithm code and can be modified by the user. Note also that |*Z*_0,0_(*ω*)|, |*Z*_0,*j*_(*ω*)| and |*Z*_0,*L*_(*ω*)| were analytically computed for each original stem dendrite using the NEURON impedance tool^64^.

### Neuron models used in the present study

To estimate the accuracy of the reduction method, we ran 50-second simulations on cell morphologies of different types, in both the reduced and detailed models. Models of 13 neurons were used in this study; their details are available in Supplementary Table 2. For each of the models, we distributed 1,250 - 10,000 synapses on their dendritic trees. Eighty percent of the synapses were excitatory, and the rest were inhibitory. The synaptic conductances were modeled using known two-state kinetic synaptic models^17^. For simplicity, we did not include synaptic facilitation or depression. All models had one type of GABAA -based inhibitory synapses and either AMPA - or AMPA+ NMDA-based excitatory synapses. The synaptic rise and decay time constants were taken from various works cited in Supplementary Table 2. When no data were available, we used the default parameters of the Blue Brain Project synaptic models^17, 53^. Inhibitory synapses were activated at 10 Hz, whereas the activation rate of the excitatory synapses was varied to generate different output firing rates in the range of 1 to 20 Hz (**Figures 2, 3, 4, 7 and Supplementary Figures 3**-**7**); the values used for each model are listed in Supplementary Table 2. In all models except Supplementary Fig. 4, synaptic activation time was randomly sampled from a homogenous Poisson process. In **Supplementary Fig. 4 a, b** the activation time was sampled from an inhomogeneous Poisson process with a time-dependent intensity λ(*t*) = *r* * sin(*t* * *f* * 2*π*) + 1, where *t* is time in seconds, r is the firing rate of the synapse and *f* is the oscillation frequency.

In Supplementary Fig. 4**c** we extracted a single Layer 5 Thick-tufted Pyramidal Cell with an early bifurcating apical tuft (L5_TTC2; gid 75586) from active blue brain microcircuit^17^. Calcium and potassium concentration were set to 1.23 mM and 5.0 mM, respectively. The synaptic activation from the microcircuit was replayed to this detailed model and also to its respective reduced model. Synaptic depression and facilitation were disabled, and the synapse time constants, which varied in the microcircuit, were set to their mean value (the decay time constant was set to 1.74 ms and 8.68 ms for AMPA and GABAA respectively; the rise time constant for GABAA was set to 4.58 ms) all other variables were as in the blue brain simulations.

### Estimating the accuracy of the reduced models

Cross-correlations were calculated between the spike trains of the detailed and the reduced models. The window size was 500 ms, and the bin size was 1 ms. The resulting cross-correlations were normalized by the number of spikes in the detailed model (**Fig. 2c**). Inter spike intervals (**ISI**) were binned in windows of 21 ms to create the ISI distribution in **Fig. 2d**.

SPIKE-synchronization measure is a parameter- and scale-free method that quantifies the degree of synchrony between two spike trains^44^. SPIKE-synchronization uses the relative number of quasi-simultaneous appearances of spikes in the spike trains. In this study, we used the Python implementation of this method^65^. To allow comparison to the literature, Supplementary Figure 3 depicts three additional metrics against which to compare the performance of the detailed and the reduced models: Trace accuracy^39^, ISI distance^44^, and Γ coincidence factor^66^.

### Comparison to other reduction algorithms

We compared *Neuron_Reduce* to two classical reduction algorithms (Supplementary Fig. 5):

#### 1. Equivalent cable using the d3/2 rule for reduction

Rall and Rinzel^32, 33^ showed that for idealized passive dendritic trees, the entire dendritic tree can be collapsed to a single equivalent cylinder that is analytically identical (from the point of view of the soma) to the detailed tree. However, neurons do not have ideal dendritic trees, mostly because dendritic terminations typically occur at different electrotonic distances from the soma. Nevertheless, it is possible to collapse any dendritic tree using a similar mapping (Rall’s “d3/2 rule”) as in the idealized tree; this will provide an “equivalent cable” (rather than an “equivalent cylinder”) with a varying diameter for the whole dendritic tree (see details in Rall et al.^48^). The electrotonic distances to the soma of synapses and non-linear dendritic mechanisms were computed in the original model and then each synapse and mechanism was mapped to the corresponding segment in the “equivalent cable” preserving the electrotonic distance to the soma as in the original tree.

#### 2. Mapping all synapses to the soma

Another recent reduction scheme was proposed where all dendritic synapses are mapped, after implementing cable filtering for each synapse, to the somatic compartment^34^. Here we used a modified version of this method. We used *Neuron_Reduce* to generate a multi-cylindrical model of the cell as in **Fig. 1b**. Then, all the synapses in the original tree were mapped to the model soma. To account for dendritic filtering, for each synapse, we multiplied the original synaptic conductance, *g_syn_*, by the steady-state voltage attenuation factor from the original dendritic location, *j*, of the synapse to the soma. Specifically,

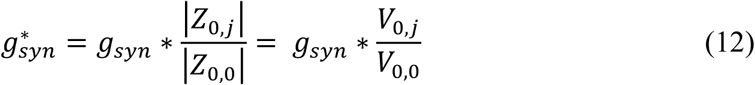

where 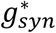 is the new synaptic weight for synapse *j* when placed at the soma of the reduced model.

### Discriminating between spatio-temporal patterns of synaptic activation

In **Figure 6**, twelve synapses, placed at 25 µm intervals, were distributed on a stretch of one basal dendrite. The peak AMPA conductance per synapse was 5 nS. In cases where the synapses also had an NMDA component, the NMDA-based peak conductance was 3.55 nS. The synapses were activated in a specific temporal order with a time delay of 3.66 ms between them. This resulted in an input velocity of 7 µm/s for the sequential IN and OUT patterns in **Fig. 6**. In addition, the temporal order of synaptic activation was randomized and scored according to the directionality index^52^, which sums the number of swaps used by the bubble sort algorithm to sort a specific temporal pattern into the IN pattern. In this measure, an IN pattern is attributed the value of 0 (no swaps) and the OUT pattern the value of 67 (67 swaps in bubble sort are required to “sort” the OUT pattern into the IN pattern^52^).

All simulations were performed using NEURON 7.4-7.7^20^ running on the Blue Brain V (BB5) supercomputer based on HPE SGI 8600 platform hosted at the Swiss National Computing Center (CSCS) in Lugano, Switzerland. Each compute node was composed of an Intel Xeon 6140 CPUs @2.3 GHz and 384GB DRAM. Analysis and simulation were conducted using Python and visualization using Matplotlib^67^

The *Neuron_Reduce* algorithm is publicly available on GitHub (http://github.com/orena1/neuron_reduce).

## Acknowledgment

We thank Gal Eliraz for her early work on the reduction method and to Mickey London for advising us along this project. This study received funding from the European Union’s Horizon 2020 Framework Program for Research and Innovation under the Specific Grant Agreement No. 785907 (Human Brain Project SGA2), the ETH domain for the Blue Brain Project (BBP), from the Gatsby Charitable Foundation and from the NIH Grant Agreement U01MH114812.

## Author contributions

I.S proposed the principle theoretical idea for the *Neuron_Reduce* scheme. I.S, O.A, E.G., and N.R. extended the original idea, planned and designed the study. O.A, E.G., and N.R implemented the *Neuron_Reduce* simulations. F.S and K.P assisted with the detailed benchmarking of *Neuron_Reduce*. M.G helped refactor the tool to increase its usability, maintainability and comprehensibility. All authors wrote the manuscript.

## Data availability

The *Neuron_Reduce* algorithm and most of the models that were used in the paper are publicly available on GitHub (http://github.com/orena1/neuron_reduce).

## Supplementary Information

**Supplementary Figure 1.**
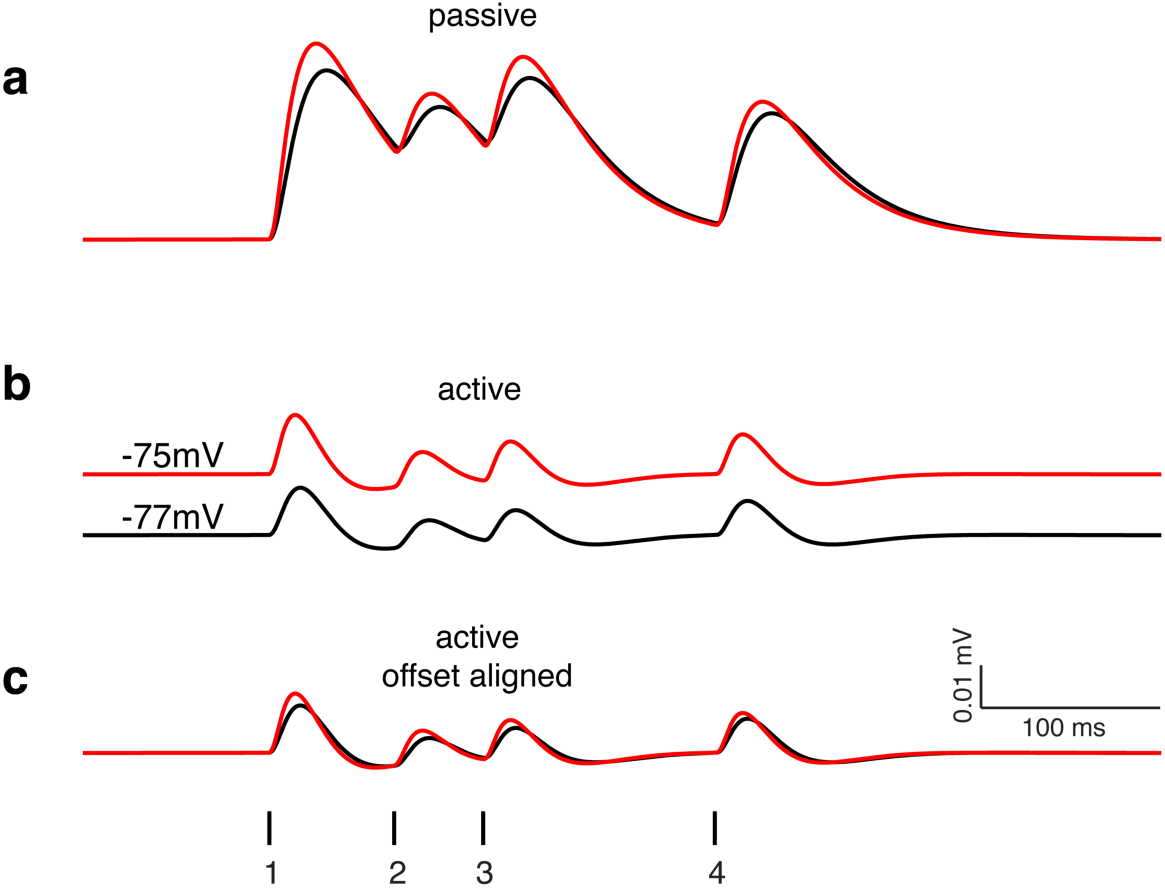
Voltage response for synaptic activation in passive and active models of L5 pyramidal cells. **a** Sequential activation of the four synapses shown in Fig. 1c in a passive model of an L5 pyramidal cell (black) and its reduced model (red), as in Fig. 1e. **b** Activation of the same synapses, but on the detailed active models of this cell (similar colors). Note that in the active model, the resting potential in the reduced model is more depolarized than in the detailed model. **c** same as in b, but with the super-positioning of the resting potential in the two models.

**Supplementary Figure 2.**
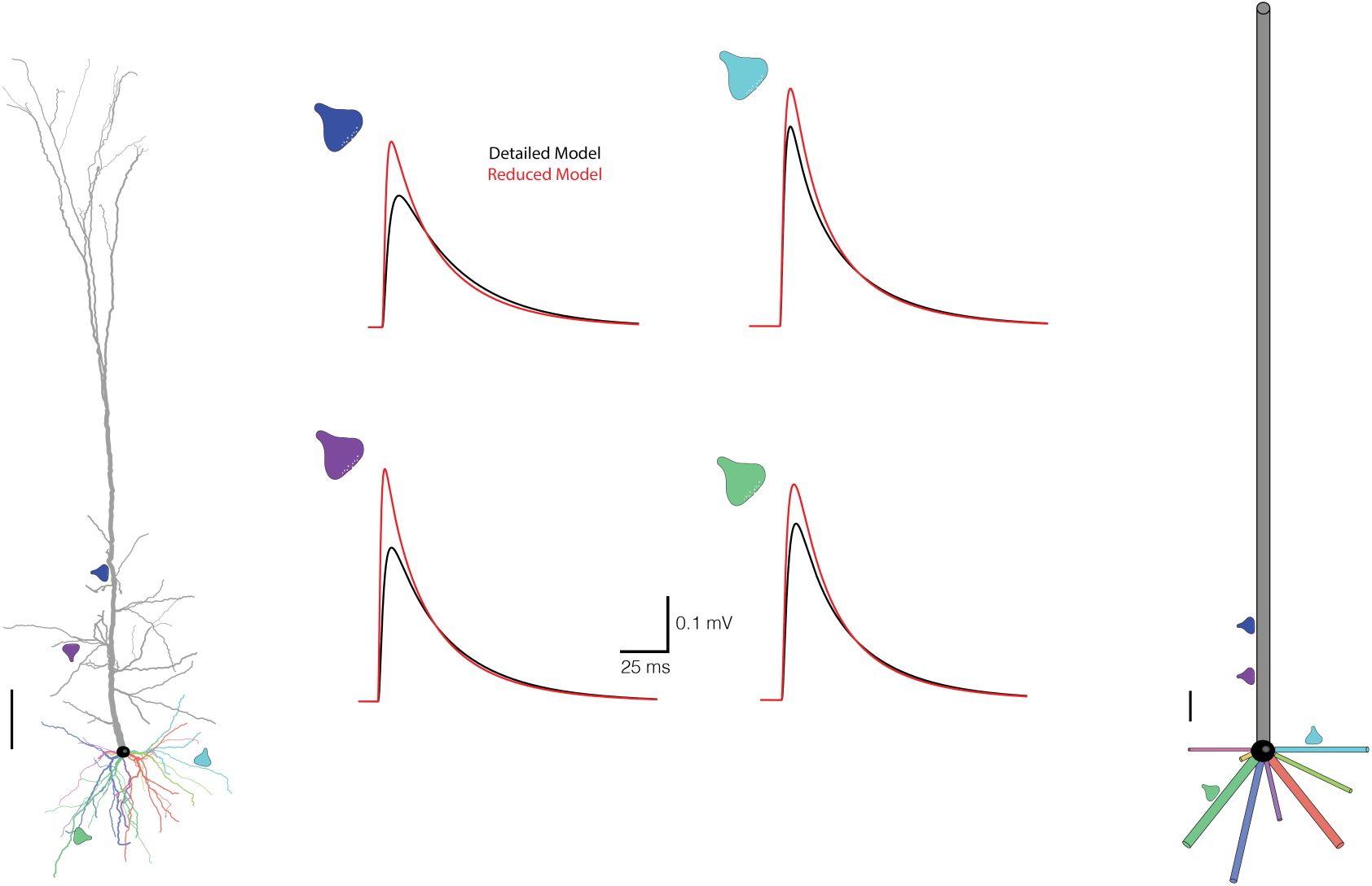
Somatic voltage response in the detailed vs. the reduced model of a layer 5 pyramidal cell. Left and right, the detailed and reduced models as in Fig. 1. Middle, EPSPs measured at the soma of the detailed (black traces) and reduced (red traces) models for the four simulated synapses. Location of synapses is shown by respective colored synapses on the tree; their EPSPs are also shown by the respective colored synapses near the voltage traces. Note that as expected from Fig. 1d, at high frequencies (e.g., at the EPSP peak) the reduced model overestimated the peak value at the soma, but the time-integrals of the EPSP in the two models remain similar (they are identical, in the passive case, since the time-integral behaves as in the steady-state case^1^). Synapse rise and decay times were 0.3 and 1.8 ms respectively, and the peak synaptic conductance was 0.8 nS. Scale bar is 100 μm in both the detailed and reduced models.

**Supplementary Figure 3.**
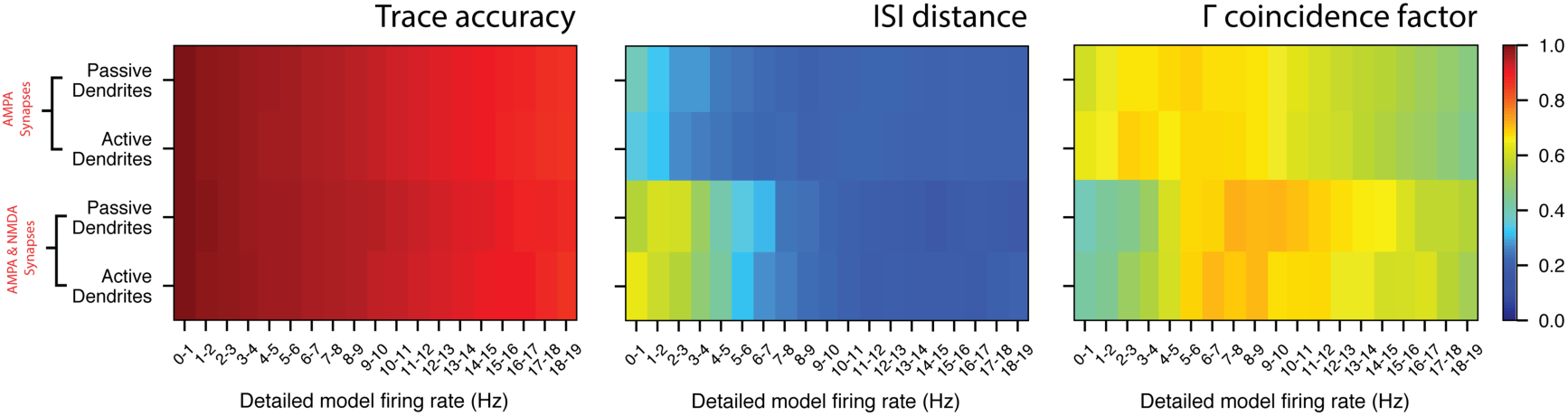
*Neuron_Reduce* performance quantified using different synchronization metrics. The performance of *Neuron_Reduce* on the L5PC shown in Fig. 2a as compared to that of the detailed model as a function of the firing rate of the detailed model, for active and passive dendrites and with/without NMDA-based synaptic conductance (same conditions as in Fig. 2g). Three different quantification methods for the quality of *Neuron_Reduce* performance are shown: Trace accuracy^2^, ISI distance^3^, and the Γ coincidence factor^4^. In both the Trace accuracy and the Γ coincidence factor, higher values represent better synchronization while in the ISI distance method, lower values represent higher synchronization.

**Supplementary Figure 4.**
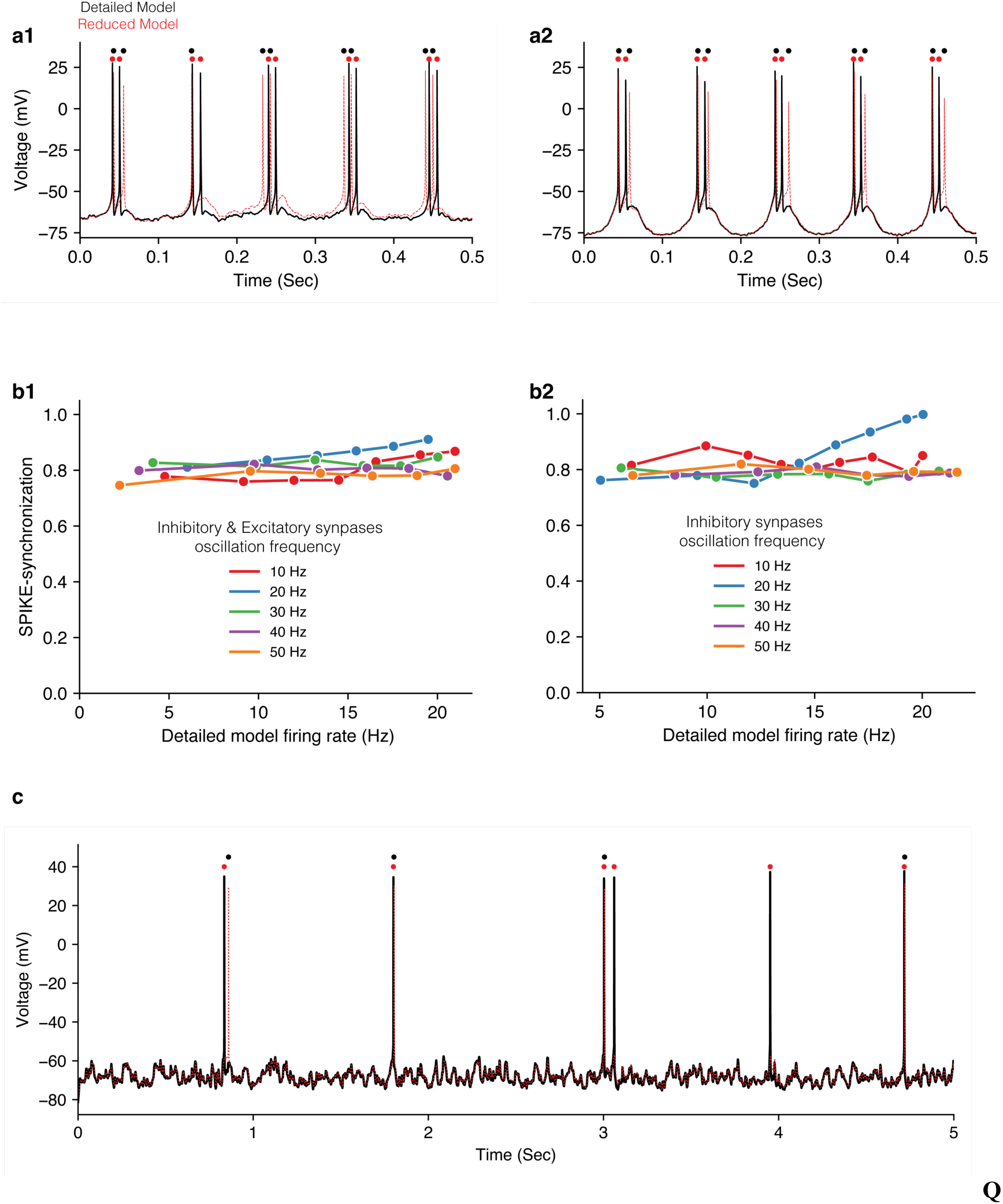
*Neuron_Reduce* performance under realistic synaptic input. **a1** Voltage dynamics at the soma of the detailed model (black trace) and the reduced model (red trace) of the L5PC model shown in Fig. 2a. The inhibitory and excitatory input was oscillating at 10 Hz; the excitatory synapses were activated at a rate of 6.7 Hz (see **Methods**). **a2** Same as in a1, but in this case only the inhibitory synapses were oscillating. **b1** SPIKE-synchronization as a function of the detailed model firing rate for different oscillating frequencies for the case where both excitatory and inhibitory synapses were oscillating. **b2** As in b1, but when only the inhibitory synapses were oscillating. **c** Voltage dynamics at the soma of the detailed model (black trace) and of the reduced model (red trace) of a Layer 5 Thick-tufted Pyramidal Cell with an early bifurcating apical tuft (L5_TTC2) extracted from active Blue Brain microcircuit^5^ (see **Methods**). The synaptic activation from the microcircuit was replayed to the detailed model and the resultant somatic voltage is shown in black. The cell was simplified using *Neuron_Reduce* and the resultant voltage trace is shown in red. SPIKE-synchronization was of 0.71.

**Supplementary Figure 5.**
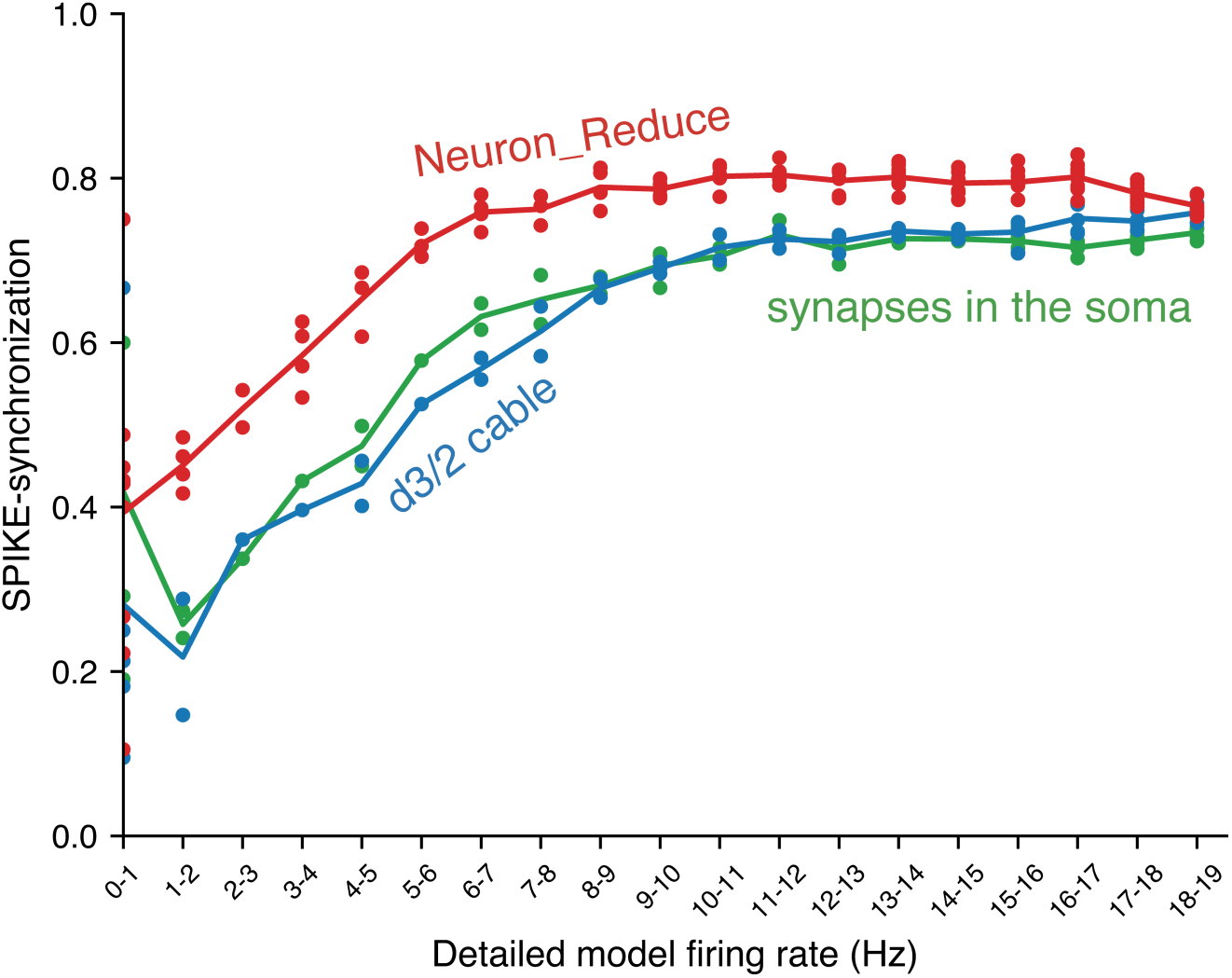
The accuracy of *Neuron_Reduce* compared to two other simplification methods. Accuracy was measured via SPIKE-synchronization as a function of the firing rate of the detailed model. Three reduction methods are compared: *Neuron_Reduce* (red), Rall’s d3/2 reduction method (“equivalent cable” approach, blue), and the mapping of all the synapses to the soma (green) see **Methods** for details on these reduction methods. Note that the accuracy of *Neuron_Reduce* is better than that of the other two methods across the entire range of model responses.

**Supplementary Figure 6.**
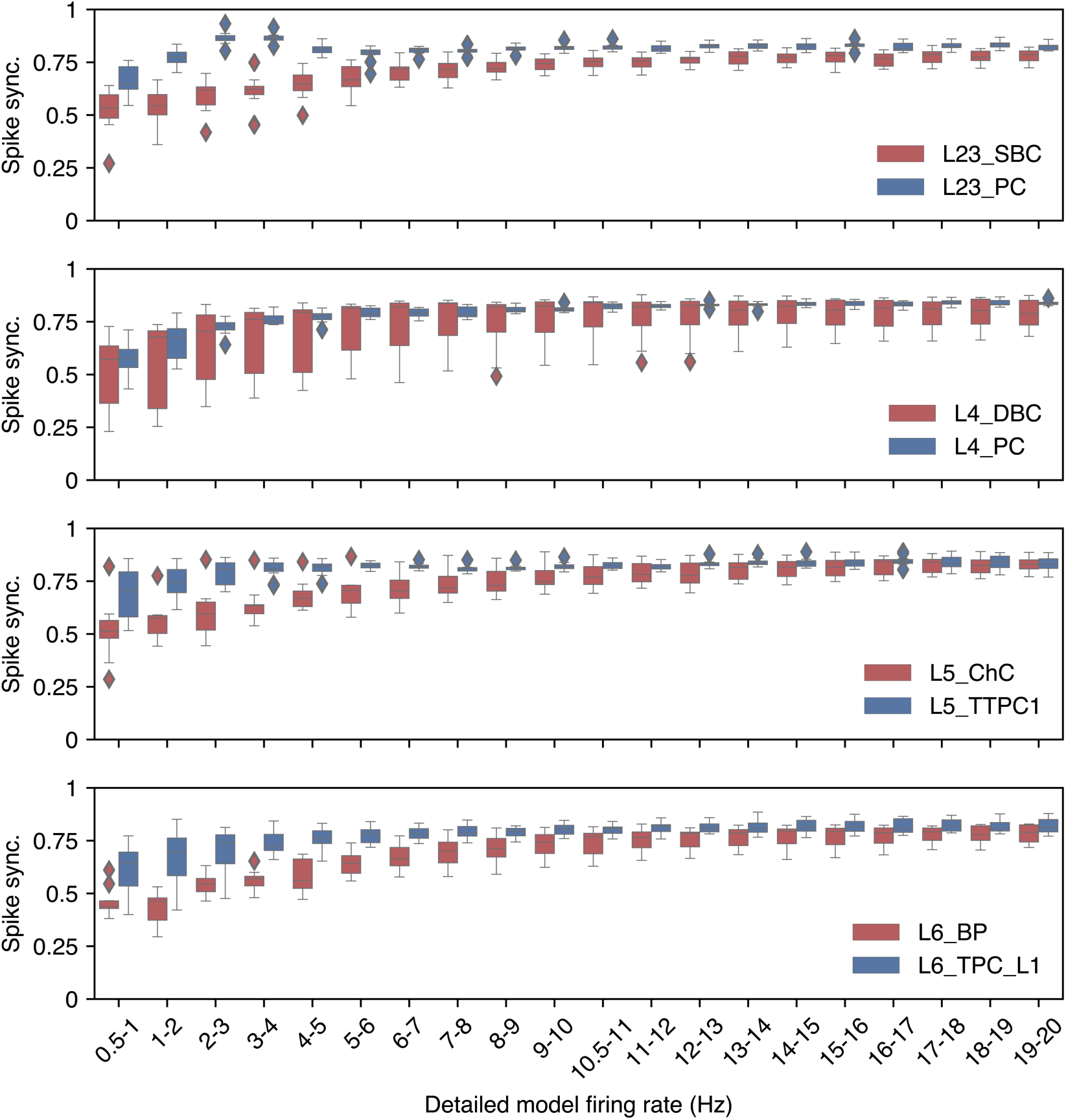
The accuracy of *Neuron_Reduce* on a large sample of neocortical neurons. Boxplot (whiskers extend to ± 1.5 interquartile range (IQR)) of Spike-synchronization measured on 8 morphological types from the Blue Brain database^5, 6^. 11 models were sampled for each type (88 cells in total), and simulations were executed for 30 seconds for each output frequency (x-axis). Each excitatory neuron model received 10,000 synapses, and each inhibitory neuron received 3,000 synapses. Synaptic input for the excitatory neurons was as for tufted PC (L6) shown in Supplementary Table 2 (5^th^ row). Synaptic input for inhibitory neurons was as for the L5 Martinotti cell shown in Supplementary Table 2 (9^th^ row). The number of compartments in the reduced model was set to 10% of the number of compartments in the detailed model when spatial discretization, ΔX, per compartment of 0.1λ resulted in a total number of compartments that is lower than 10% of the number of compartments in the detailed model (See Supplementary Table 1). SBC – Small Basket Cell; DBC – Double Bouquet Cell; ChC – Chandelier Cell; BP – Bipolar Cell; PC – Pyramidal Cell; TTPC1 – Thick-tufted Pyramidal Cell with a late bifurcating apical tuft; TPC_L1 – Tufted Pyramidal Cell with apical dendrites terminating in layer 1.

**Supplementary Figure 7.**
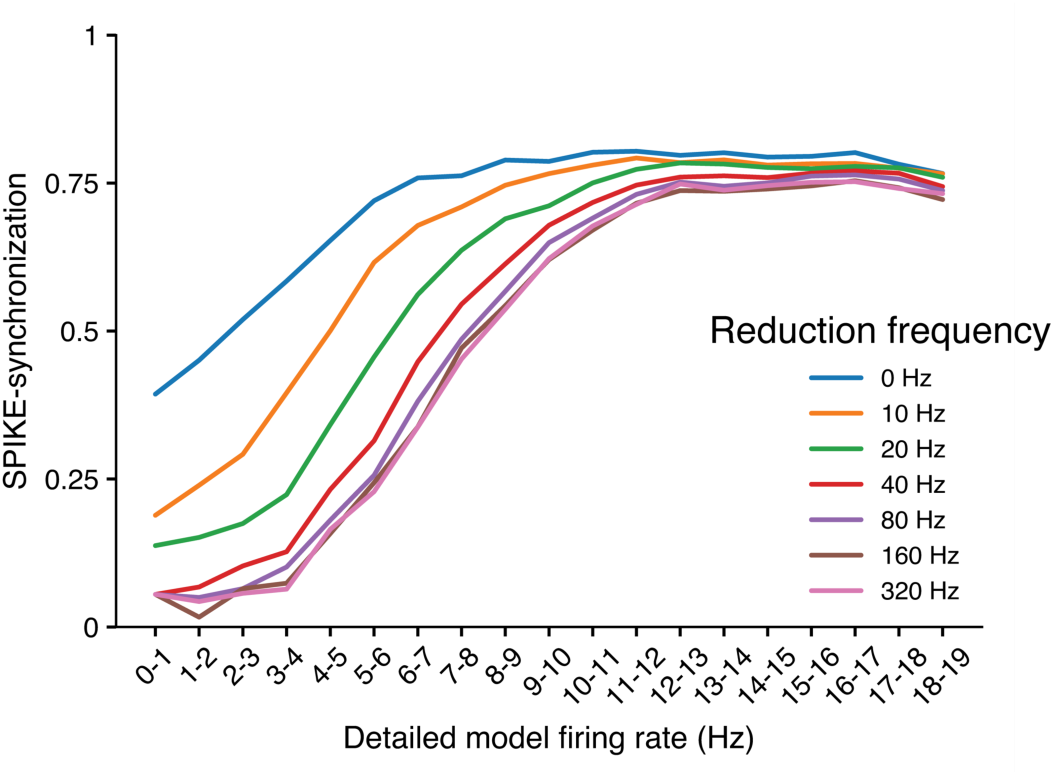
Reduction performance ω values. Spike synchronization as a function of the spike firing in the detailed model for different frequencies, ω, used in Neuron_Reduce (Eqs. (1)-(11)). Model used as in Fig. 2f for the case denoted by the orange line. Note that the case of ω = 0 outperforms all other cases.

**Supplementary Table 1.** Enhanced simulation speeds in the reduced models versus the detailed models of a L5 pyramidal cell. The modeled cell is shown in Fig. 1; simulations were performed with 10,000 synapses; simulation time was 50 seconds, excitatory and inhibitory synapse firing rate was 5 and 10 Hz, respectively (see Fig. 2 and 3). Total run time for the detailed model is shown in the first row, first column of the shaded region. In the second row, the detailed dendritic morphology (consisting of 642 compartments) was mapped to its respective multipolar cylindrical representation consisting of only 50 compartments (as in Fig. 1b), whereas the total number of synaptic point processes (10,000) was kept as in the detailed model (1.39 folds reduction in run-time). In the third row, the detailed morphology remained untouched but synapses located on the same compartment were merged into a single point process (∼2 point processes/compartment, one representing all E-synapses and the other all I-synapses located on this compartment). This resulted in 1234 synapses and a 2.9-fold reduction in run-time. In the bottom row, we used the full *Neuron_Reduce* algorithm, in which both the morphology reduction and the synaptic merging algorithms were implemented (50 compartments and 94 synaptic point processes), resulting in a 45-fold reduction in total simulation run-time (see also Fig. 3a). Columns 4 and 5 shed light on the underlying reasons for the speed-up: the runtime is dominated by two contributions, a) the number of compartments and b) the number of synapses. In brackets, we also list the resulting runtimes (Please note: since the total runtime has a third contribution from solving the linear system of the branched tree (not shown), the runtimes given in brackets do not add up to the total runtime). In the detailed model, the contributions from the compartments and synapses are 1/3 and 2/3, respectively. In the cases of only morphological reduction or only synaptic merging, the speed-up factor accordingly is limited by that contribution’s weight in the sum. Only when reducing both at the same time, the full speed-up can be realized. The small runtime difference for 50 compartments in the morphological reduction vs. *Neuron_Reduce* case is attributed to more memory overhead in the case of the morphological reduction with a much larger number of synapses. Note that these calculations are based on 10 repetitions for each condition, on a cluster with Intel Xeon 6140 CPUs and NEURON version 7.4 (see **Methods**). For each single cell, we used a single core on each node allocated exclusively to avoid side effects from other running processes.

| Reduction steps | Total run-time (s) | Run time speed-up factor | Number of compartments<br>(run-time, s) | Number of synapses<br>(run-time, s) |
| --- | --- | --- | --- | --- |
| Detailed model | 2,906 | 1 | 642 (796.5) | 10,000 (1,942.6) |
| Morphological reduction | 2,036 | 1.4 | 50 (55.7) | 10,000 (1,891.4) |
| Synaptic merging | 1005.5 | 2.9 | 642 (762.8) | 1234 (154.5) |
| *Neuron_Reduce* | 68.7 | 42 | 50 (46.3) | 94 (8.5) |

**Supplementary Table 2.**
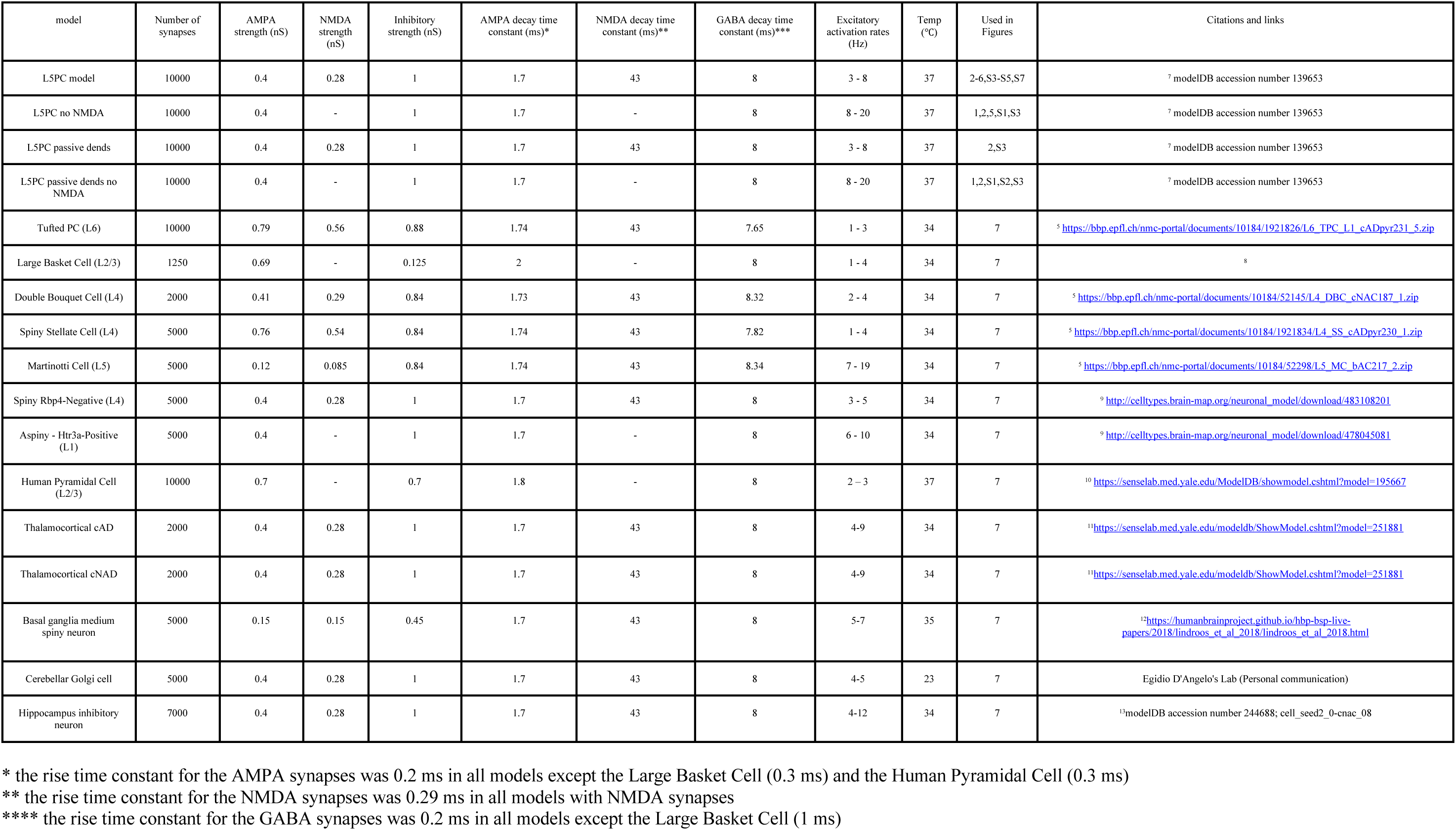
Neuron models and synaptic parameters used in this article.

